# *Agrobacterium*-mediated *Cuscuta campestris* transformation as a tool for understanding plant-plant interactions

**DOI:** 10.1101/2024.02.23.581736

**Authors:** Supral Adhikari, Asha Mudalige, Lydia Phillips, Hyeyoung Lee, Vivian Bernal-Galeano, Hope Gruszewski, James H. Westwood, So-Yon Park

**Author notes:** These authors contributed equally to this article (Supral Adhikari, Asha Mudalige).

## Abstract

*Cuscuta campestris*, a stem parasitic plant, has served as a valuable model plant for the exploration of plant-plant interactions and molecular trafficking. However, a major barrier to *C. campestris* research is that a method to generate stable transgenic plants has not yet been developed. Here, we describe the development of a *Cuscuta* transformation protocol using various reporter genes (GFP, GUS, or RUBY) and morphogenic genes (*CcWUS2* and *CcGRF/GIF*), ultimately leading to a robust protocol for *Agrobacterium*-mediated *C. campestris* transformation. The stably transformed and regenerated RUBY *C. campestris* plants produced haustoria, the signature organ of parasitic plants, and these were functional in forming host attachments. The locations of T-DNA integration in the parasite genome were confirmed through TAIL-PCR. Transformed *C. campestris* also produced flowers and transgenic seeds exhibiting betalain pigment, providing proof of germline transmission of the RUBY transgene. Furthermore, the RUBY reporter is not only a useful selectable marker for the *Agrobacterium*-mediated transformation, but also provides insight into the movement of molecules from *C. campestris* to the host during parasitism. Thus, the protocol for transformation of *C. campestris* reported here overcomes a major obstacle to *Cuscuta* research and opens new possibilities for studying parasitic plants and their interactions with hosts.

## Introduction

*Cuscuta* spp. are obligate stem parasitic plants that infect a broad range of host plants (Dawson *et al*., 1994). The thread-like plant consists of leafless, twining stems that can attach to aerial parts of herbaceous plants, woody shrubs, and even trees. The parasitic invasion of *Cuscuta* begins by coiling around the stem of its host plant and initiating points of growth toward the host. The epidermal cells of these adhesive disk structures secrete a bioadhesive substance that anchors the parasite to the host (Vaughn, 2002; Galloway *et al*., 2020). Next, a multicellular feeding structure termed the haustorium develops from the parasite stem and breaches the host tissues using a combination of mechanical pressure and enzymatic digestion of host cell walls, ultimately resulting in vascular connections with the host (Nagar *et al*., 1984; Johnsen *et al*., 2015; Kaiser *et al*., 2015).

*Cuscuta* spp. are among the most agriculturally destructive parasitic weeds, as well as interesting subjects for studies of plant-plant interaction. The economic impact is due to their wide range of potential host species, which includes many dicotyledonous crops and ornamental plants (Lanini & Kogan, 2005), as well as their habit of extracting water and nutrients that the host needs for productivity. For example, *C. campestris* reduced tomato fresh weight by 98% in a greenhouse study (Goldwasser *et al*., 2012), and *C. indecora* decreased alfalfa yield by 57% (Cudney *et al*., 1992). Although cultural, chemical, and mechanical control methods have been developed to suppress the growth of *Cuscuta*, there is currently no efficient method to eliminate established *Cuscuta* without also causing damage to their hosts (Dechassa & Regassa, 2021).

*Cuscuta* spp. are useful for plant biology because their parasitic lifestyle provides insights into the mechanisms by which plants detect and interact with other plants. Examples of this include *Cuscuta* detection of neighboring plants by volatile signals (Runyon *et al*., 2006), *Cuscuta*-to-host bidirectional mobility of mRNAs, small RNAs, and proteins (Kim *et al*., 2014; Shahid *et al*., 2018; Liu *et al*., 2020; Park *et al*., 2022), and elucidation of host resistance mechanisms against parasites (Hegenauer *et al*., 2016; Hegenauer *et al*., 2020). Therefore, both applied and basic plant sciences would benefit from greater ability to investigate the molecular mechanisms underlying interactions between *Cuscuta* spp. and *their* hosts.

Plant genetic transformation is an essential tool for functional genomic studies as it enables the manipulation of gene expression through the creation of knockout or overexpression mutant lines. This approach has been used for selected species of parasitic plants in the family Orobanchaceae to understand the mechanisms by which they develop and interact with their hosts. Methods for the transient and stable transformation of these root parasitic plants have been published for *Triphysaria versicolor, Phtheirospermum japonicum*, and *Phelipanche aegyptiaca* (Tomilov *et al*., 2007; Fernández-Aparicio *et al*., 2011; Ishida *et al*., 2011; Bandaranayake & Yoder, 2018).

As for *Cuscuta* transformation, several efforts have led to partial success. Regeneration of shoots from callus of *C. reflexa* (Srivastava & Dwivedi, 2001) has been reported, and transformation was accomplished in cultured *C. trifolii* (Borsics *et al*., 2002) and *C. europeae* (Švubová & Blehová, 2013). Transformation of haustorial structures was recently accomplished in *C. reflexa* (Lachner *et al*., 2020). However, despite these attempts with diverse *Cuscuta* species, the inability to regenerate a functional *Cuscuta* plant from transformed explant tissues has blocked progress toward achieving stably transformed *Cuscuta*.

The failure to regenerate plants from transgenic callus is a common bottleneck in plant transformation, but progress has come from the use of genes involved in morphogenesis, which promote the formation of plant organs in callus of plant species that are recalcitrant to transformation or regeneration (Lee & Wang, 2023). For example, recent reports have shown improved regeneration and transformation efficiency in maize (*Zea maize*) with morphogenic transcription factors such as *WUSCHEL* (*WUS*) and *BABY BOOM* (*BBM*). *WUS*, previously known to play a vital role in maintaining the stem cell population, was also found to enhance embryogenesis in *Arabidopsis* (Zuo *et al*., 2002). When used with a transformation recalcitrant maize line, *BBM*-*WUS* enhanced transformation efficiency by over 40% (Lowe *et al*., 2016). In another approach, a chimera consisting of *GROWTH-REGULATING FACTOR* (*GRF*) and its co-factor *GRF-INTERACTING FACTOR* (*GIF*) was shown to promote transformation and regeneration efficiency in wheat (*Triticum aestivum*) and citrus (*Citrus sinensis Osb. × Poncirus trifoliata L. Raf.*) (Debernardi *et al*., 2020). *GRF* and *GIF* are transcription factors regulating growth and development of plants, and their ability to enhance the efficiency of transformation has been extended to maize (*Zea mays*), watermelon (*Citrullus lanatus*), and lettuce (*Lactuca spp*). The combination of *GRF-GIF* with *BBM* improved the transformation efficiency of maize seedlings from 26% to 37% (Chen *et al*., 2022). Furthermore, in watermelon, overexpression of the *GRF-GIF* chimeric gene resulted in higher transformation efficiency than control conditions (Feng *et al*., 2021). The *GRF-GIF* chimera with mutated miRNA396 improved the regeneration and shooting efficiency in lettuce (Bull *et al*., 2023).

Here we describe a transformation protocol for *C. campestris* that was developed through exploration of several different reporter genes and morphogenic genes. Although *Cuscuta* species have been broadly regarded as recalcitrant to transformation, the technique readily generates transgenic *C. campestris* plants that show normal coiling and haustorium formation, and ultimately produce flowers and seeds. This approach unlocks new possibilities for conducting functional studies on *Cuscuta* and facilitates investigations into understanding of plant-plant interactions.

## Materials and methods

### *Cuscuta* seed scarification and germination

*C. campestris* seedlings were attached to 3-week-old beets (Detroit Dark Red, *Beta vulgaris*) and grown in a greenhouse with a temperature of 25°C - 30°C and a day/night length of 16/8 hours. The use of beet as a nursery host for *C. campestris* is a coincidence that has nothing to do with the use of the RUBY marker; beet has a low growth profile, plentiful petioles, and high susceptibility, so it is an excellent host for the parasite. Three months after the inoculation, *Cuscuta* seeds were collected and cleaned. To promote germination, clean seeds were scarified as described previously (Bernal-Galeano *et al*., 2022). Scarified seeds were rinsed with autoclaved water three times, sterilized with 3% sodium hypochlorite for 15 minutes, washed with autoclaved water five times and incubated at 4°C overnight. For seed germination, 100 *Cuscuta* seeds were placed on a petri dish with a wet Whatman’s 3MM paper and incubated at 28°C without light for two days. The petri dish containing *Cuscuta* seedlings was placed to a growth chamber set to 25 °C with 16 hours of light and 8 hours of darkness using Lithonia Lighting Fluorescent Luminaire bulbs (Model: C 2 32 120 GESB) that provided 300-400 lux of illumination and incubated for one day to be 3-day-old seedlings.

### *Agrobacterium-*mediated *Cuscuta* transformation

To develop the *Agrobacterium*-mediated *Cuscuta* transformation protocol, we tested three different conditions including optimization of media growth hormone concentrations, simplified inoculation process with RUBY, and stable generation of *Cuscuta* transgenic lines.

First protocol - *Optimization of media growth hormone concentrations*: A single colony of *A. tumefaciens* EHA105 containing the *35S:GUS-pMAS:GFP* (Supp Figure 1A) was inoculated into 1-2 mL of liquid MGL medium and incubated at 29°C with shaking at 200 rpm for 48 hours. The bacterial culture was then spread on a solid MGL plate and incubated overnight at 29°C. The resulting creamy bacterial culture was used for inoculating the explant (Supp Figure 1B). Three-day-old seedlings (Supp Figure 1C), as described in the seed germination section, were infected with *Agrobacterium* using the streak infection method from the previous protocol (Kanchanamala & Bandaranayake, 2019). The 3-day-old seedling’s apical part without meristem was cut from each plant (Supp Figure 1D-E) (Furuhashi *et al*., 2012) and dipped in the creamy bacteria culture for 3-5 seconds (Supp Figure 1F). The explants were placed on co-cultivation media (CCM) containing 3 mg/L Naphthalene Acetic Acid (NAA) (N0640, Sigma) and 1 mg/L Benzylaminopurine (BAP) (B3274, Sigma) for co-cultivation at 15°C for 7 days with 16 hours of light and 8 hours of darkness condition (Supp Figure 1G and Supp Table 1). Subsequently, the infected explants (Supp Figure 1G) were transferred to MMS media containing antibiotic cefotaxime (250 mg/L) (C-104-5, GoldBio), cytokinins, and auxins (Supp Figure 1I and Supp Table 1). We tested NAA at concentrations of 0 and 0.5 mg/L, BAP at 5, 7.5, 10, and 25 mg/L, Thidiazuron (TDZ) (P6186, Sigma) at 0.05, 0.1, 0.3, and 0.5 mg/L and MMS media without any growth hormones. We designed a factorial experiment to observe the ideal plant growth hormone combination for better transformation efficiency.

Second protocol - *Simplified inoculation process with RUBY*: To simplify the inoculation process, we developed a bulk chopping method that is less precise but quicker than carefully dissecting out the shoot apical region. The laborious dissecting method was termed the “cut” technique, and the more random method was termed the “bulk chop” technique. We compared the transformation efficiencies between the cut and bulk chop methods for preparing *Cuscuta* segments. Using a sterilized surgical blade, a large mass of 3-day-old seedlings from 200 seeds was chopped into small pieces to make around 5mm segments, which greatly facilitated the tissue preparation step and produced a large number of potentially transformable sections at once. Also, instead of a construct with GUS and GFP reporters, we used a construct containing *35S:RUBY* (He *et al*., 2020). *A. tumefaciens* EHA105 harboring the construct *35S:RUBY* was strecked on a LB plate and incubated at 28 °C for two days. A single colony was cultured in 5mL liquid media and grown in a shaker at 28 °C for 16 hours. This culture was subsequently subcultured in 50mL of LB liquid media and incubated for another 16 hours at 28 °C. When the OD600 reached 1.0 to 1.2, the cells were harvested and resuspended in liquid MMS media without a growth hormone. Infected explants were placed on CCM for 5 days and then transferred to CIM containing Cefotaxime (250 mg/L) and BAP (5 mg/L) (Supp Table 1). Transformed tissues expressing RUBY were counted around a month after inoculation.

Third protocol - *Stable generation of Cuscuta transgenic lines*: Three-day-old *Cuscuta* seedlings were bulk-chopped into small pieces as described in the second protocol. The bacteria were prepared following a procedure similar to the second protocol. After cocultivation with *A. tumefaciens* EHA105 solutions, *Cuscuta* seedlings were uniformly distributed and incubated on co-cultivation media for five days (Supp Table 2). After the co-cultivation stage, infected seedlings were transferred to Callus Induction Media (CIM) containing 5 mg/L BAP and 250 mg/L cefotaxime (Supp Table 2). For the GUS staining assay, calli were harvested 30-35 days after infection. To generate stably transformed *Cuscuta*, calli were incubated on CIM for at least 8 weeks, shoot induction media (SIM) for at least 8 weeks, and shoot elongation media (SEM) for 3-4 weeks (Supp Table 2). Media with explants were replaced with new media every two weeks.

The co-cultivation, callus induction, shoot induction, and shoot elongation processes for the first, second, and third protocols were conducted under the following conditions: 25 °C with 16 hours of light and 8 hours of darkness using Lithonia Lighting Fluorescent Luminaire bulbs (Model: C 2 32 120 GESB), providing 300-400 lux of illumination.

### GUS staining and GFP observation

For the GUS staining, *Cuscuta* calli were incubated in X-Gluc solution (G1281C2, GoldBio) for up to 24 hours and then destained in 95 % Ethanol (Park *et al*., 2022). Images were captured using Leica Thunder imager M205 FA model stereomicroscopy system.

Fluorescent signals were detected with both Leica Thunder imager M205 FA model stereomicroscopy system and Leica SP8 spectral confocal microscope. All microscopy for GUS and fluorescence expression was conducted at the Advanced Light Microscopy Core (https://research.missouri.edu/advanced-light-microscopy), University of Missouri. GFP and RFP expression patterns were imaged with the following setting (excitation/emission): GFP at 488 nm / 510 nm and RFP at 561 nm / 584nm. RFP was used as a negative control.

### Vibratome sectioning and imaging

*Cuscuta* haustoria expressing RUBY or WT *Cuscuta* grown on tomato (*Solanum lycopersicum* var. Rutgers) or *Arabidopsis* Col-0 for two weeks were harvested and embedded in 5 % Agarose (Park *et al*., 2022). Tissue sectioning was performed using a Leica series Vibratome 3000 Plus. Longitudinal and transverse sections were cut with thickness ranging from 250µm to 400µm, depending on the softness of the tissue. Sections were stained by 0.02% toluidine blue (T3260, Sigma) (Pradhan Mitra & Loqué, 2014). For imaging, sections were placed on microscope slides and imaged by Leica M205 FA automated stereo microscope.

### Thermal asymmetric interlaced (TAIL)-PCR

To confirm T-DNA insertions in transgenic *Cuscuta*, genomic DNAs of transgenic *Cuscuta 35S:RUBY* No. #9a and No. #18a were extracted using Cetyltrimethylammonium-bromide (CTAB) (Sigma), and TAIL PCRs were conducted using insertion-specific primers (RB1 and RB2) and arbitrary degenerate primers (AD primers) using primary and secondary PCR conditions (Supp Tables 3 and 4). Specific bands from amplification with AD and RB2 primers were extracted and sequenced. The presence of T-DNA insertions in transgenic RUBY *Cuscuta* plants was verified by aligning the sequencing results with a *Cuscuta campestris* genome sequence (Vogel et al., 2018).

### Vector constructions

*35S:GUS-pMAS:GFP* (Supp Figure 1A) was modified by replacing the 35S promoter with the MAS promoter from *pGFPGUSPlus* (Plasmid #64401, Addgene) (He *et al*., 2020) because *pMAS:GFP* showed stronger expression than *35S:GFP* (Mudalige *et al*., under preparation).

**Figure 1.**
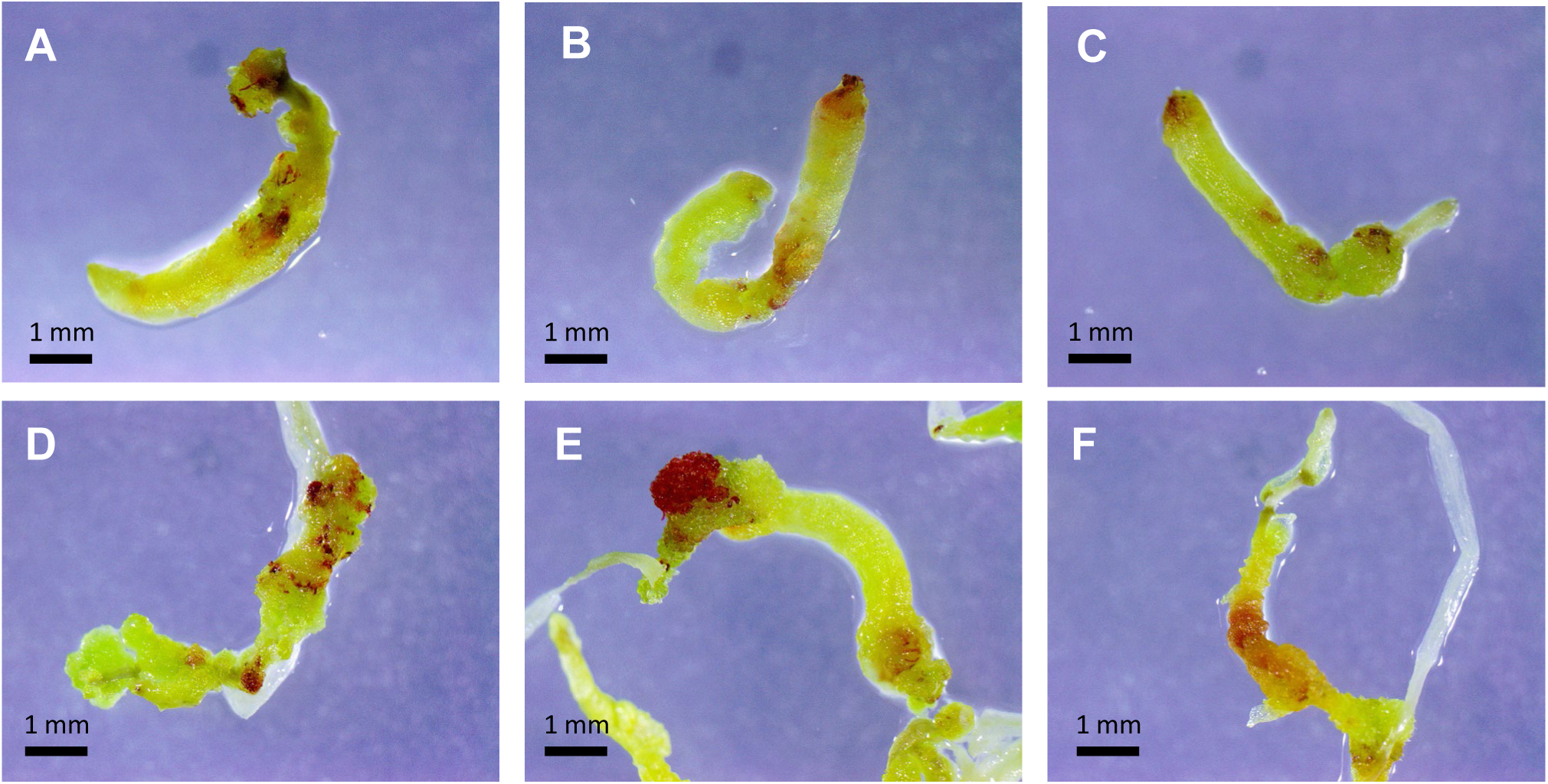
RUBY expression in explants. RUBY expression in cut explants (A, B, C) and bulk chopped explants (D, E, F) approximately 30 days after inoculation.

*CcGRF* and *CcGIF* were identified from the *C. campestris* genome (Vogel *et al*., 2018) via sequence alignments and phylogenetic trees including wheat homologs (Debernardi *et al*., 2020), performed in the Geneious Prime 2022.2.2 software. The selected *CcGRF* homolog Cc010427 and *CcGIF* homolog Cc021394 were cloned and sequenced using gene specific primers (Supp Table 5). The confirmed clones were ligated to make a *CcGRF/CcGIF* fusion via the Golden Gate method using NEB’s NEBridge kit and re-cloned into Pcr8GWTOPO vector (Invitrogen).

*CcWUS2* was synthesized by Azenta life science (www.azenta.com) and re-cloned into Pcr8GWTOPO vector.

*CcWUS2* and *CcGRF/GIF* in entry clone pCR8GWTOPO were incubated with LR clonase and pMDC139 to generate destination vectors with genes of interest (*35S:CcWUS2-GUS* and *35S:CcGRF/CcGIF-GUS*). *35S:GUS-MAS:GFP*, *35S:CcWUS2-GUS*, and *35S:CcGRF/GIF-GUS* were transformed into *A. tumefaciens* EHA105 strain by the freeze-thaw transformation method (Weigel & Glazebrook, 2006).

## Results

### Test of hormones and media composition for *C. campestris* transformed calli

To establish the transformation protocol for *C. campestris*, we initially tested various concentrations of NAA and cytokinin (BAP/TDZ) in a medium designed for both co-cultivation and callus induction (Supp Table 1) using the first protocol described in the material and method section (Supp Figure 1). A negative control, consisting of the same medium without growth hormones, was also employed. Approximately thirty days after inoculation with *A. tumefaciens* EHA105-*35S:GUS-pMAS:GFP* (Supp Figure 1A), transformation efficiency was assessed by GFP expression using a stereomicroscope (Supp Figure 2A-D). Cells expressing GFP (Supp Figure 2A and 2D) but not RFP (Supp Figure 2B) were counted as transformed tissue (Supp Table 6). Additionally, a confocal microscope revealed that GFP was located in the nucleus and cytoplasm, which is typical GFP localization (white arrows, Supp Figure 2E). The merged image of GFP, RFP, and bright field demonstrated GFP-specific expression in a month-old explant (Supp Figure 2D, 2H, and 2I).

Using the first transformation method and a GFP reporter construct, we compared transformation efficiencies between different treatments (Supp Figure 3 and Supp Table 6). The statistical analysis indicated a significantly higher GFP expression in the media containing plant growth regulators compared to that without hormones (Supp Figure 3). *Cuscuta* explants cultivated on CIM media with BAP or TDZ without NAA exhibited significantly higher GFP expression than those on CIM media with NAA or the no hormone media, indicating that the presence of cytokinin hormones had a positive effect while the presence of NAA had a negative impact on efficiency. However, there was no significant difference among treatments with different concentrations of BAP or TDZ. Thus, we concluded that the presence of BAP or TDZ without NAA in the CIM culture media significantly enhanced the *C. campestris* transformation efficiency. For subsequent experiments, we used 5 mg/L BAP to induce *Cuscuta* callus in CIM media.

### Comparison of the transformation efficiency of explants prepared with the cut versus the bulk chop method using the RUBY reporter

While approximately 50% of *Cuscuta* calli expressed GFP 30 days after *A. tumefaciens* inoculation and cultivated on 5mg/L BAP media (Supp Figure 2), the preparation of individual meristems through the cut method (Supp Figure 1D) is labor-intensive, as it involves the individual handling of each seedling. We tested a simpler bulk chop method that produced a large amount of *Cuscuta* segments for potential transformation. In this comparison, both cut and bulk chop samples were infected with *A. tumefaciens* EHA105-*35S:RUBY* and incubated on CIM for 30 days (Supp Table 1). RUBY encodes enzymes for the synthesis of betalain and is useful for visually monitoring transformation events without staining or special equipment. Calli derived from the cut method exhibited 100% RUBY expression, whereas calli from the bulk chop method displayed relatively lower RUBY expression (65%) (Supp Table 7). Previously we found thatthe shoot apical part of *Cuscuta* generated more callus and transformable tissue than other parts (Mudalige et al., under preparation). Thus, we hypothesize that the bulk chop method had lower RUBY expression than the cut method because the non-transformable tissues including root or shoot tips were included with the transformable tissues incubated on the media. In addition, calli expressing RUBY from the cut method (Figure 1A-C) did not exhibit any phenotypic differences when compared to calli from the bulk chop method (Figure 1D-F). As RUBY has been used as a visual selection marker in other plants (Yu *et al*., 2023), we decided to employ it as a visual selection marker during the early co-cultivation stage, eliminating the need for a selection reagent.

### *Agrobacterium-*mediated *Cuscuta* transformation and regeneration of transgenic plants

To conduct the *Agrobacterium*-mediated *Cuscuta* transformation, three-day-old seedlings from 200 *C. campestris* seeds were chopped and infected with *A. tumefaciens* EHA105-*35S:RUBY* as described in the third protocol, the method section. *Agrobacterium* infected segments were incubated on the co-cultivation media for five days (Figure 2A). Subsequently, 100 selected *Cuscuta* segments expressing strong RUBY were divided among two CIM media plates, each containing 50 explants per plate (Figure 2B). Explants were incubated in CIM media for a minimum of 8 weeks before being transferred to shoot induction media (Figure 2C-D). Larger and darker red calli were obtained after another 8 weeks of incubation on the shoot induction media (Figure 2E-F) and then transferred to shoot elongation media when the red calli reached a size of 1 cm (Figure 2 G-H). Calli grown on the shoot elongation media started to produce shoots in two weeks (Figure 2I).

**Figure 2.**
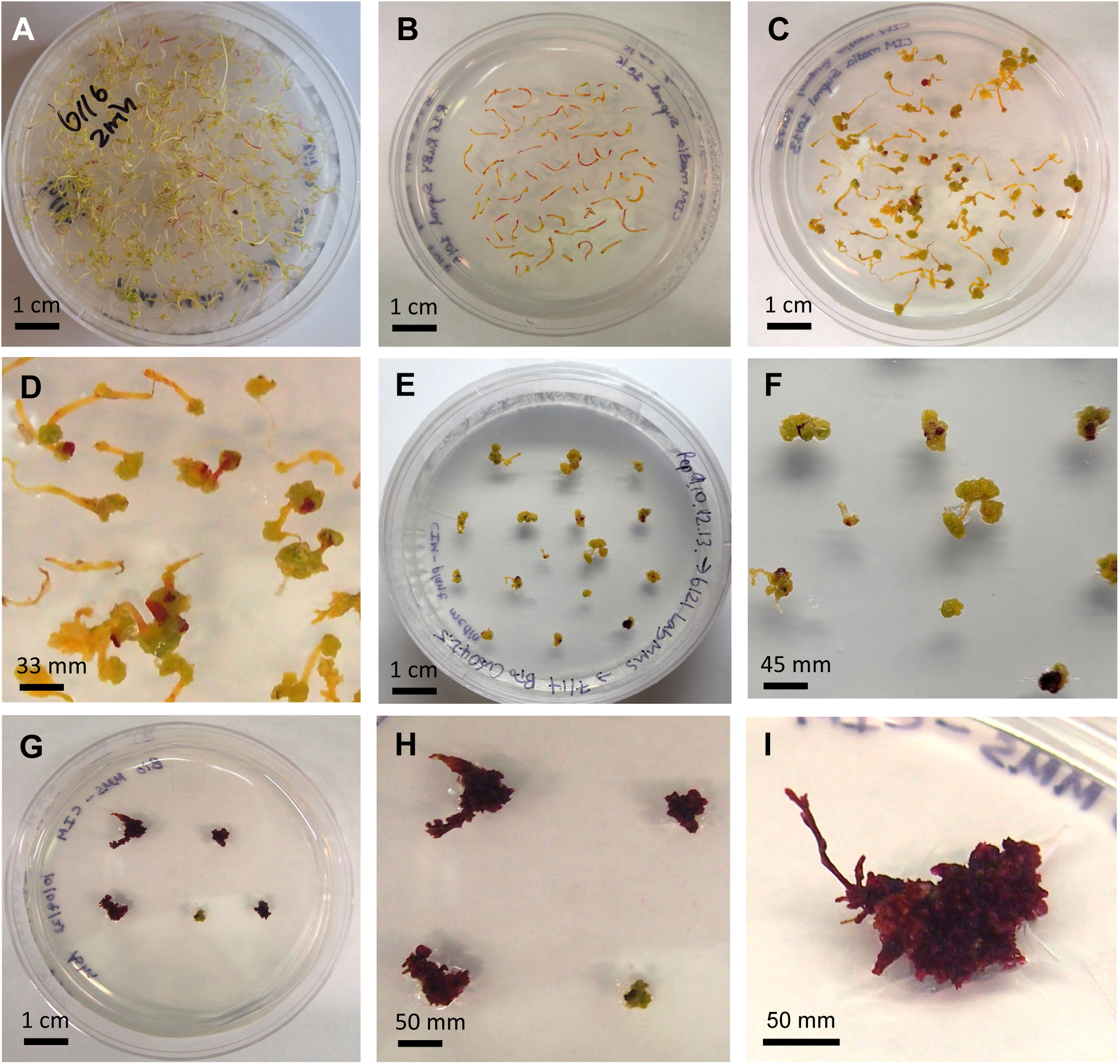
Process of *Agrobacterium*-mediated *Cuscuta* transformation using *A. tumefacient* EHA105 containing *35S:RUBY*. *Cuscuta* seedlings were infected by EHA105-*35S:RUBY* and incubated on CCM for 5 days (A). *Cuscuta* segments expressing RUBY were transferred on CIM from CCM (B). One-month-old RUBY calli and were grown on CIM (C-D) and SIM (E-F). Calli expressing RUBY were selected and transferred to SEM (G-H) and produced shoots (I). (D), (F), and (H) are zoomed images of (C), (E), and (G), respectively.

During the callus induction stage, the number of visible red spots on *A. tumefaciens* EHA105-*35S:RUBY* infected *Cuscuta* segments were counted on 5, 15, 25, 35, and 45 days after infection to track the transformation efficiency. Since we selected 100 *Cuscuta* segments expressing RUBY from 200 *Cuscuta* seeds to callus induction media after 5-day co-cultivation on each replicate, transformed calli represented 100% on the fifth day of inoculation. The number of transformed calli slowly declined to 93.2%, 86.0%, 76.7 %, and 62.1% on the 15, 25, 35, and 45 days after inoculation, respectively (Figure 3A and Supp Table 8).

**Figure 3.**
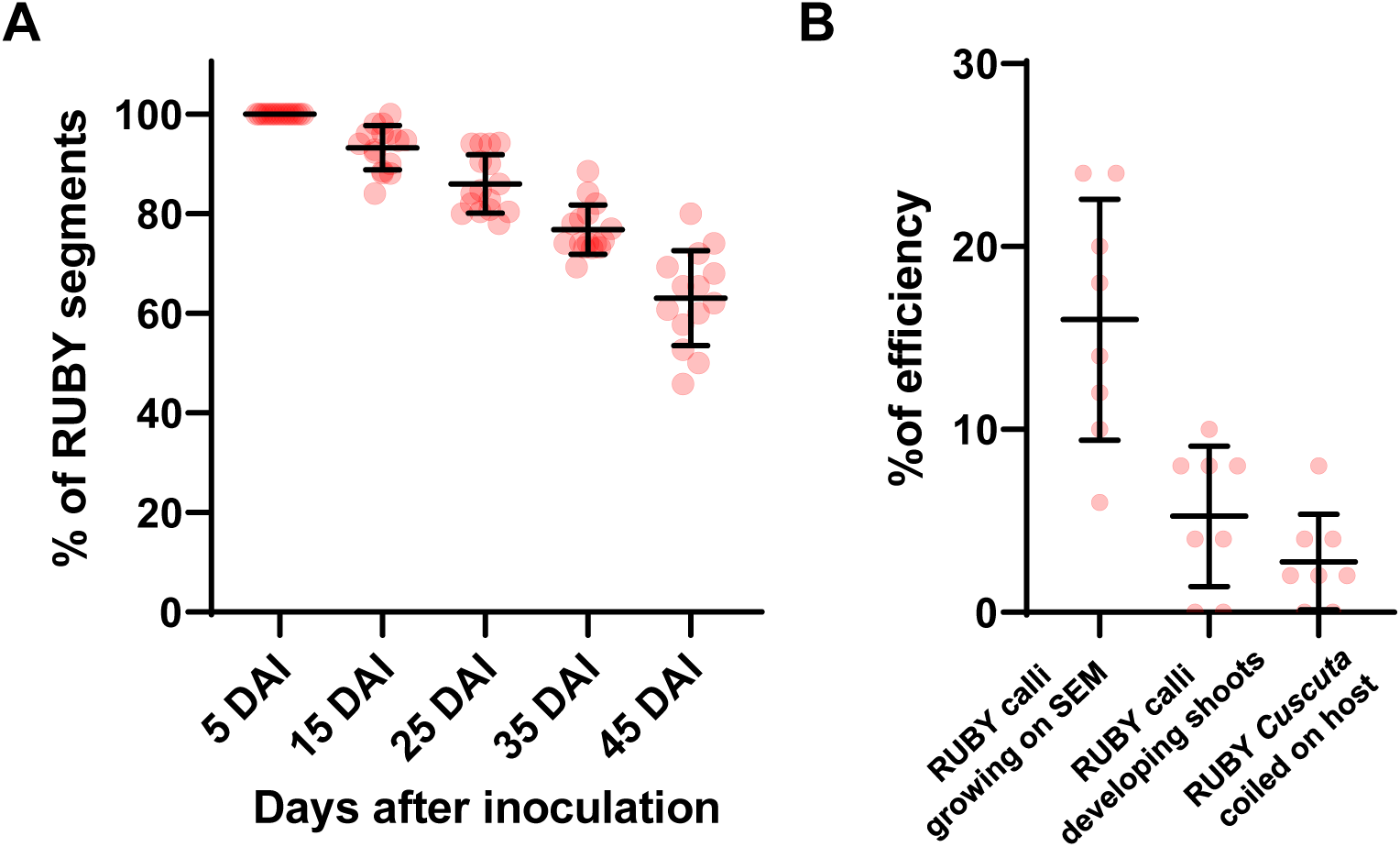
*C. campestris* transformation efficiency with the RUBY reporter gene. (A) Transformation rate on various days following inoculation. Sixteen independent replicates were performed. (B) The percentage of RUBY calli growing on SEM, RUBY calli producing shoots, and RUBY *Cuscuta* coiled on a live host was quantified. Eight independent replicates were performed. Data are presented as the mean ± standard deviation. Each replicate had 100 explants from 200 *C. campestris* seeds and split into two CIM plates, each containing 50 explants per plate.

Further declines in the number of stably transformed calli occurred over subsequent growth stages. Percentages of RUBY calli growing on SEM, RUBY calli developing shoots, and RUBY *Cuscuta* coiled on host plants were 16%, 5.3%, and 2.8%, respectively (Figure 3B and Supp Table 9).

### Successful parasitism of transgenic RUBY *Cuscuta*

Once the RUBY *Cuscuta* shoots reached a size of 3 cm on the shoot elongation media, calli with RUBY *Cuscuta* shoots were moved to magenta boxes containing the shoot elongation media (Figure 4A). To observe the characteristic coiling behavior of *Cuscuta*, a toothpick was inserted into the media to serve as a substitute host. Within a day of introducing the toothpick, RUBY *Cuscuta* coiled (Figure 4B) and produced haustoria (Figure 4C). We also introduced RUBY *Cuscuta* stems to live host plants. Within 24 hours post-inoculation, the RUBY *Cuscuta* stems exhibited coiling and initiated haustoria development on a 1-week-old tomato (Figure 4D). All host plants, both tomatoes (Figure 4E) and *Arabidopsis* (Figure 4F) were successfully parasitized by the RUBY *Cuscuta*. Approximately 2 months after transfer to beet hosts, RUBY *Cuscuta* produced flowers (Figure 4G-H) and seeds (Figure 4I).

**Figure 4.**
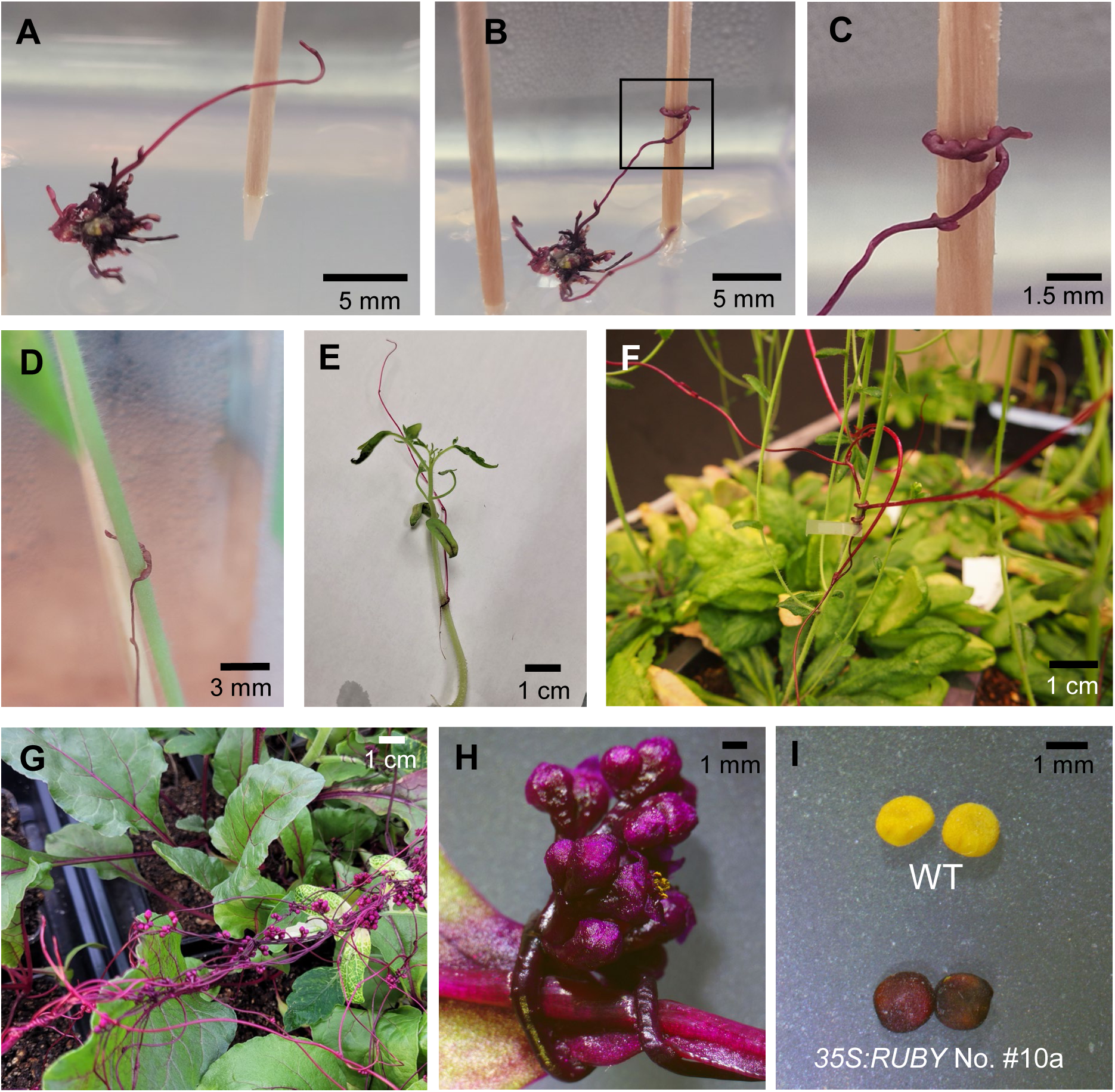
Transgenic RUBY *Cuscuta* attaching to non-living and living hosts. A shoot from transgenic callus (A) coiled on a toothpick (B-C). (C) Zoomed-in image of the black box in (B) showing haustoria formation. RUBY *Cuscuta* coiled on (D) and parasitizing (E) the stem of a 2-week-old tomato. (F) RUBY *Cuscuta* was successfully grown on *Arabidopsis*. Transgenic RUBY *Cuscuta* produced flowers on beets (G-H) and seeds (I).

Seeds from a T0 transgenic *Cuscuta* (*35S:RUBY* No. #10a1) were germinated to analyze RUBY and non-RUBY segregation ratios. Transgenic *Cuscuta 35S:RUBY* No. #11 exhibited similar germination efficiency compared to wild-type *Cuscuta* with RUBY and non-RUBY segregation ratio of 2.33 : 1 ± 0.58 (Supp Table 10). The transgenic seedlings showed distinct betalain pigments throughout their entire tissues (Supp Figure 4). This suggests that the RUBY transgene from T0 transgenic *Cuscuta* is stably inherited by the next generation through *Agrobacterium*-mediated *Cuscuta* transformation.

### Event-specific detections of transgenic lines

From 8 attempts of the *Agrobacterium*-mediated *Cuscuta* transformation, 11 transgenic events were generated and successfully coiled on host plants (Supp Table 9). To validate the T-DNA insertions, gDNA was extracted from 2 mature transgenic *Cuscuta* (No. #9a and #18a), and TAIL-PCRs using right border (RB) primers located in the 35S promoter of the *35S:RUBY* were conducted (Figure 5A). After sequencing the amplicons, results revealed that both transgenes lacked the RB sequences (gray box, Figure 5A). Furthermore, the transgene of the first event contained a complete 35S promoter, whereas the transgene from the second event had a truncated 35S promoter. By aligning TAIL-PCR sequences with *C. campestris* genome data, the integration locations of transgenes of *35S:RUBY* No. #9a and *35S:RUBY* No. #18a were located on *C. campestris* Scaffold 32B and *C. campestris* Scaffold 104, respectively (Figure 5B and Supp Figure 5). Thus, this result indicates that the transgenic events were independent transgenic plants.

**Figure 5.**
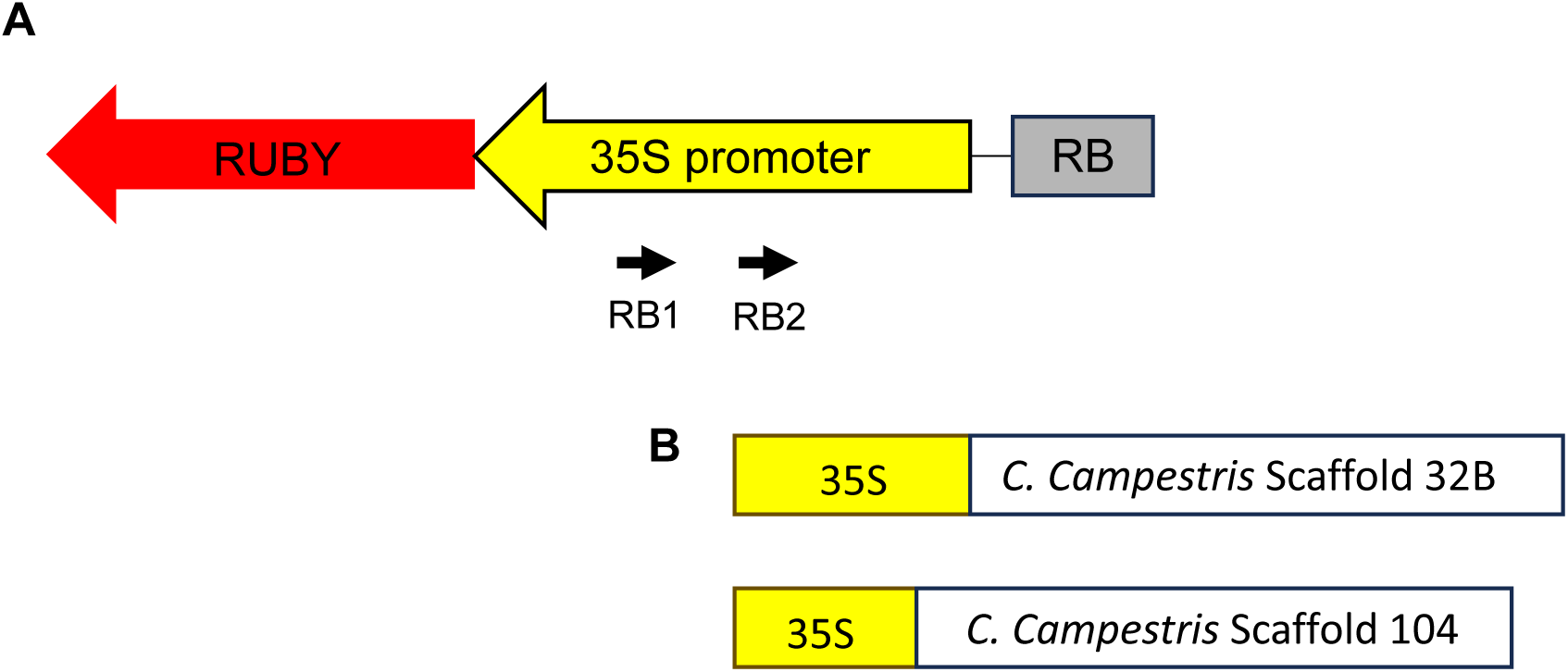
Alignment of sequences from TAIL PCRs and the *35S:RUBY* construct. (A) Diagram of 35S promoter and RUBY near RB region in *35S:RUBY*. Red arrow, yellow arrow, and gray box are RUBY, 35S promoter, and right border, respectively. Black arrows (RB1 and RB2) are primers used for the TAIL PCRs. (B) Schematic designs of T-DNA sequences and insertion points into the *C. campestris* genome. Yellow boxes indicate 35S promoter of *35S:RUBY* and white boxes are *C. campestris* genome sequences.

### RUBY pigments as markers of haustorial presence and activity in host stems

One advantage of having a transgenic parasite is the opportunity to explore haustorial development and function. Given that RUBY is an excellent visual marker, transgenic *C. campestris* was studied in interactions with tomato seedlings and *Arabidopsis* floral stems (Figure 6 and Supp Figure 6). The negative control tissues from tomato and *Arabidopsis* did not exhibit any pigment accumulation inside stems (Supp Figure 6A - 6B). WT *C. campestris* is shown coiled around a tomato stem with the haustoria invading the host tissue to establish vascular connections (Figure 6A - 6B). Cells of the haustoria are easily identifiable near their base (Figure 6B asterisk) but are difficult to distinguish from host cells near the haustorial tip. When RUBY *Cuscuta* coiled on the host stem but prior to penetration, the red pigment was not detected in host stems or vascular bundles (Figure 6C - 6D). Invasive growth of the haustorium is clear in the host stem during the early haustorium development (Figure 6E – 6F). Mature haustoria are visible within the host root (Figure 6G - 6H). Interestingly, we observed diffuse RUBY pigments beyond the area where haustoria cells were clearly detected (Figure 6G and white circles of 6H). Some searching hyphae cells were distinctly visible (black arrows, Figure 6H), and stand in contrast to the more diffuse pigment. The diffuse pattern of RUBY pigment was also observed when RUBY *Cuscuta* was grown on *Arabidopsis* Col-0 plants. After staining tissues with toluidine blue, we found that RUBY pigments were mainly detected on cortex and phloem of host *Arabidopsis* stem (Figures 6I-6K). RUBY was even detected in the outside surface of the host stem near haustoria (white arrows of Supp Figures 6D and 6E) and regions away from *Cuscuta* haustoria (white circles of Figures 6I and 6L and white arrows of Supp Figures 6F and 6G). However, *Arabidopsis* with WT *Cuscuta* did not exhibit any pigment accumulation on the surface (Supp Figure 6C) or inside the stem (Supp Figure 6F). The mechanism of pigment spread, whether it represents movement of the pigment itself or some combination of RUBY mRNA/protein, will be a point for future research, but illustrates the potential for using transgenic *Cuscuta* for investigating parasite-host communication.

**Figure 6.**
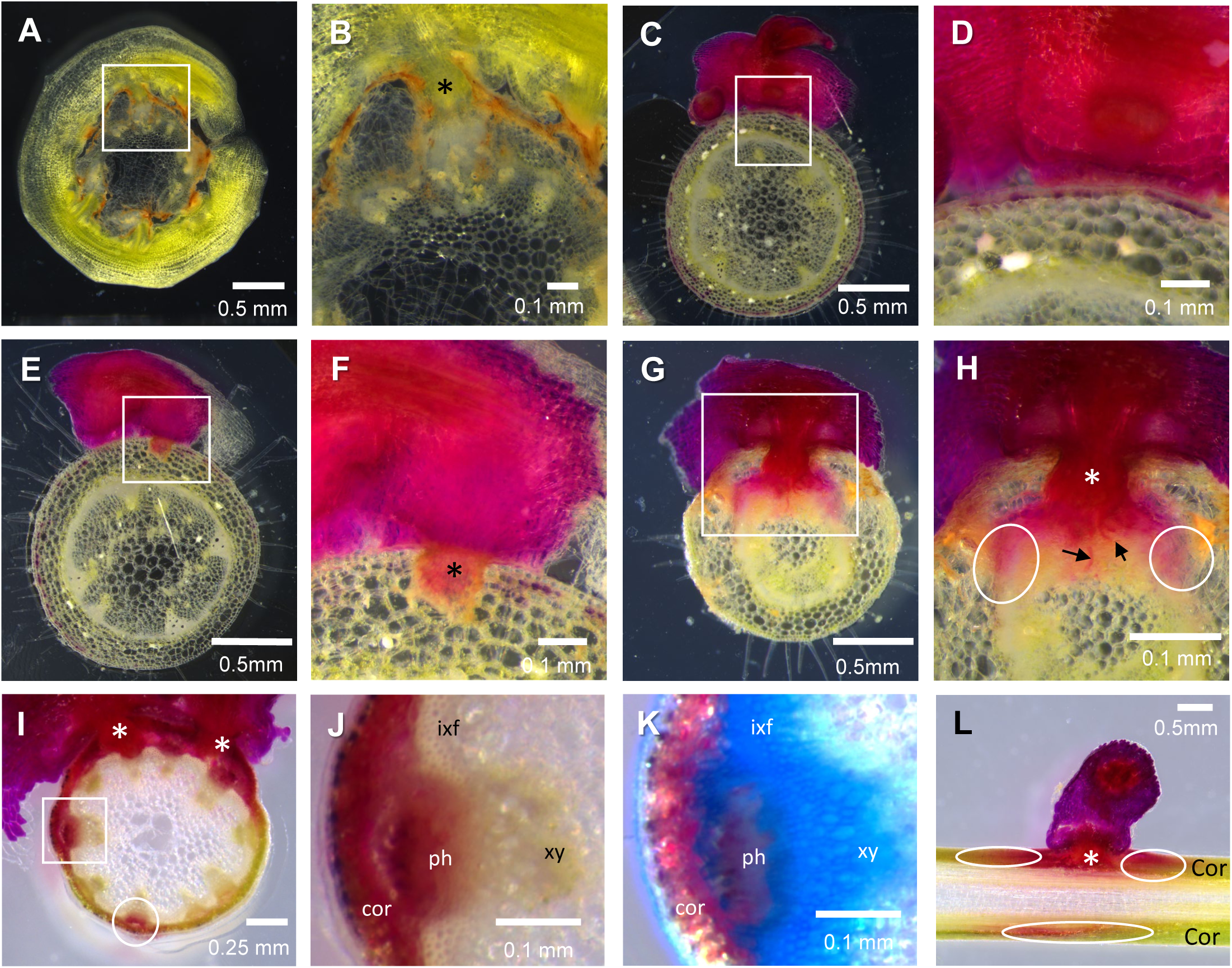
Cross sections of *Cuscuta* haustoria on tomato or *Arabidopsis* stems. Haustoria of WT *C. campestris* successfully penetrated tomato stems and established a connection with the host vascular bundle (A-B). RUBY *Cuscuta* coiling around a host tomato stem but failing to penetrate (C-D). RUBY *Cuscuta* successfully attaching to tomato is shown at early (E-F) and mature (G-H) haustoria development stages. (B, D, F, and H): Zoomed-in images of the white box region of corresponding images in (A, C, E, and G). RUBY *Cuscuta* penetrated and developed haustoria on *Arabidopsis* Col-0 stem (I). Images of the white box in (I) before staining (J) and after toluidine staining (K). Vertical section of *Cuscuta* haustorium on *Arabidopsis* Col-0 (L). Asterisks (*), white circles, and black arrows indicate the presence of *Cuscuta* haustorium, area of diffused RUBY pigment, and searching hyphae, respectively. xy, xylem; ixf, xylem fibres; ph, phloem; cor, cortex.

### Enhanced transformation efficiency using *CcGRF/GIF Cuscuta*

Morphogenic genes have been reported to enhance plant transformation efficiency (Lowe *et al*., 2016; Chen *et al*., 2022), so we tested whether *C. campestris WUS2* or *C. campestris GRF/GIF* can promote transformation efficiency. Instead of the RUBY reporter, we decided to use the GUS reporter gene to eliminate potential tyrosine deficiency phenotype caused by the RUBY reporter (Lee *et al*., 2023). Three-day-old *C. campestris* seedlings (Figure 7A) were bulk chopped and infected by EHA105 with *GUS*, *CcWUS2-GUS,* and *CcGRF/GIF-GUS* (Figure 7B). After the co-cultivation, 50 infected meristem-containing-segments were placed on CIM media (Figure 7C). Around 35 days of incubation on CIM, *Cuscuta* segments infected by EHA105 with *GUS*, *CcWUS2-GUS,* and *CcGRF/GIF-GUS* were harvested and stained (Figure 7D-F). Segments expressing GUS were quantified and classified by sizes of the GUS expressing sectors, such as no GUS, less than 0.2 mm GUS, between 0.2 to 2 mm GUS, and larger than 2 mm GUS, aiming to distinguish the extent of GUS expression in a segment (Supp Figure 7).

**Figure 7.**
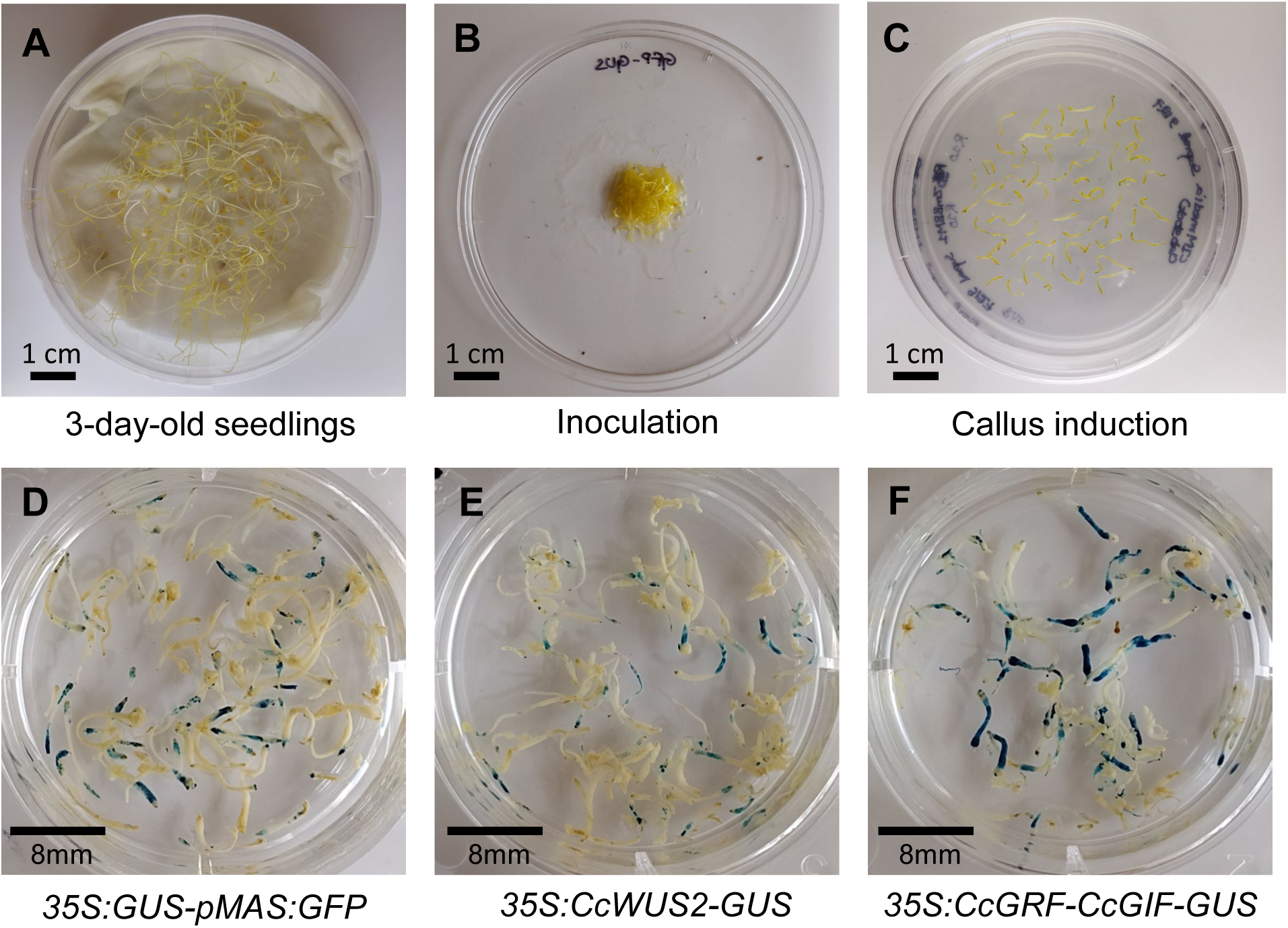
*Agrobacterium*-mediated *Cuscuta* transformation using chopped seedlings. Three-day-old seedlings (A) were chopped (B) and inoculated with *Agrobacterium tumefaciens* strain EHA105 harboring *35S:GUS-pMAS:GFP.* (C) Infected segments were incubated on cocultivation media. (D-F) Approximately 35 days after inoculation, segments infected by *35S:GUS-pMAS:GFP, 35S:CcWUS2-GUS* and *35S: CcGRF-CcGIF-GUS* were stained with X-gluc solution.

In terms of presence and absence of GUS expression, the transformation efficiencies using *GUS*, *CcWUS2-GUS,* and *CcGRF/GIF-GUS* were 78.8%, 51.7%, and 74.8%, respectively (Figure 8A). Notably, the transformation efficiency associated with *CcWUS2-GUS* was significantly lower than *GUS* or *CcGRF/GIF-GUS.* In contrast, there was no significant difference in transformation efficiency between GUS and *CcGRF/GIF-GUS*. Data from categorized segments indicated that *Cuscuta* segments transformed with *GUS* alone showed a significantly higher proportion of expressing tissues in the below 0.2 mm category compared to those transformed with morphogenic gene fusions *CcWUS2-GUS* and *CcGRF/GIF-GUS* (Figure 8B). *Cuscuta* with *CcGRF/GIF-GUS* had a significantly higher percentage in the over 2mm group than *GUS* only. These findings suggest that *GUS* alone and *CcGRF/GIF-GUS* had similar levels of transformation efficiency based on the presence or absence of GUS expression. However, *CcGRF/GIF-GUS* significantly enhanced regeneration efficiency during the callus induction stage, leading to more significant GUS expression compared to *GUS* alone.

**Figure 8.**
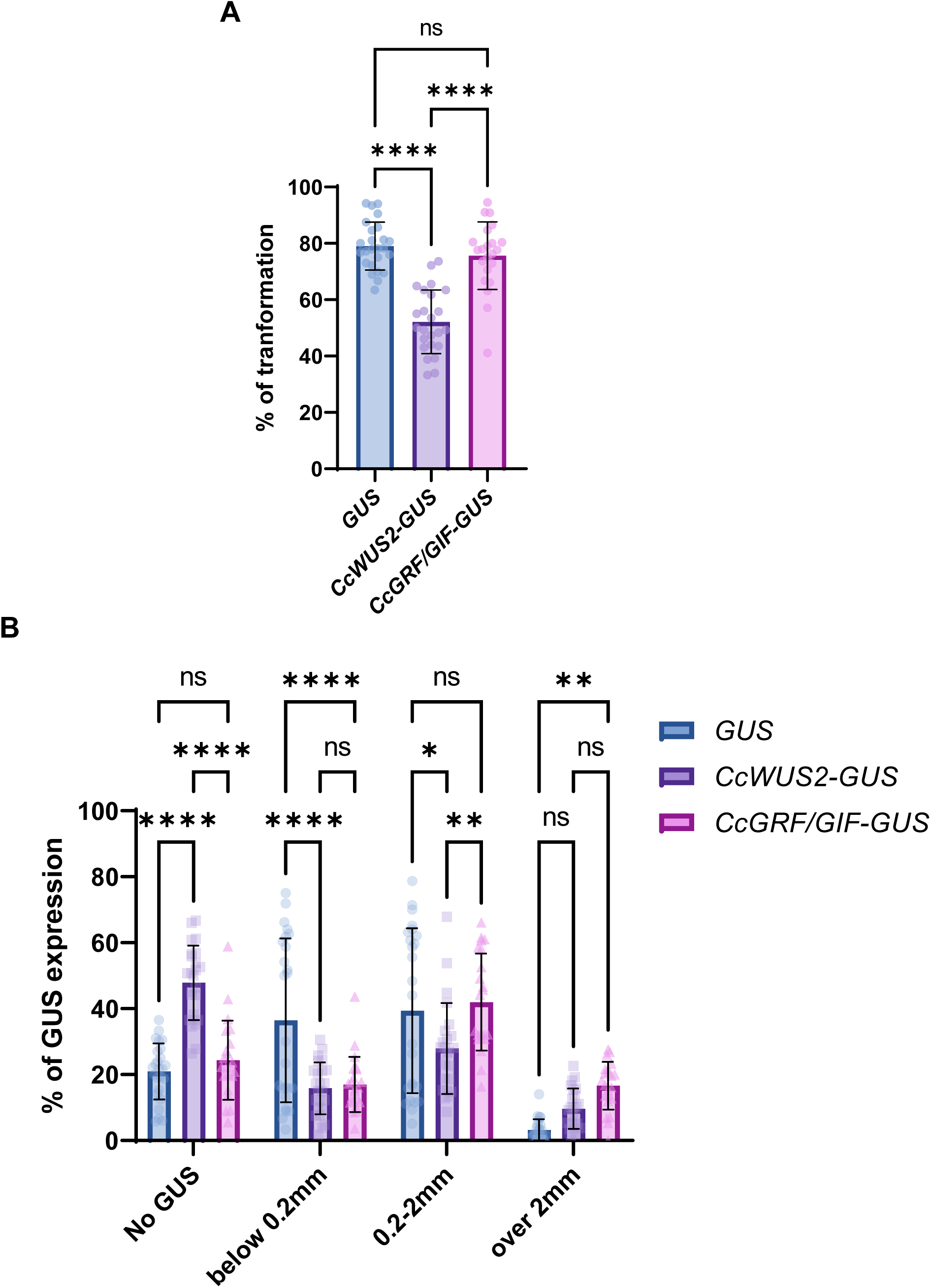
Effect of *CcWUS2* and *CcGRF/GIF* on *Agrobacterium-*mediated *Cuscuta* transformation. (A) The transformation efficiencies of *GUS* (blue bar), *CcWUS2-GUS* (purple bar), and *CcGRF/GIF-GUS* (pink bar) were compared. (B) GUS expressions were classified into four groups according to the size of GUS expression. Thirty independent replicates were performed. Data are presented as the mean ± standard deviation.

## Discussion

The lack of a protocol for transformation and regeneration of *Cuscuta* has hindered progress in studying parasite gene functions and the mechanisms of *Cuscuta*-host interactions. Despite previous studies demonstrating transformation of *Cuscuta* tissues (Borsics *et al*., 2002; Švubová & Blehová, 2013; Lachner *et al*., 2020), none have succeeded in regenerating stably transformed *Cuscuta* plants that can be used to study the full range of parasite biology. Here we describe a method for the relatively fast and efficient stable transformation of *C. campestris* plants that behave as normal parasites, producing haustoria, flowers, and seeds (Figure 4). The process hinges on *A. tumefaciens*-mediated transformation with appropriate media for inducing callus and shoots. We estimate that the process took a minimum of 5 months and resulted in a rate of 2.7% recovery of transformed plants from 100 explants expressing RUBY selected during the CIM step. While this transformation efficiency may not be high compared to easily transformable plants like tobacco, it stands out as remarkably successful for a transformation recalcitrant non-model species.

An important innovation in the transformation process was the use of the recently developed RUBY reporter (He *et al*., 2020). RUBY encodes enzymes for synthesizing betalain from tyrosine and is useful for visually monitoring transformation events without using fluorescence microscopy or a chemical selection process. However, recent studies reported that RUBY may cause low regeneration efficiency and poor fertility during seed development due to the lack of the essential amino acid tyrosine. For example, maize expressing a strong RUBY reporter had growth retardation (Lee *et al*., 2023). While we are mindful that the RUBY reporter could be detrimental to *Cuscuta*, evidence to date suggests that transgenic RUBY *Cuscuta* shows normal, though somewhat slower, growth compared to non-transgenic plants. But importantly, the vivid RUBY reporter proved to be an invaluable asset in the development of the protocol. Its strong visible color and non-reliance on toxic selection reagents make it difficult to replace with other selectable markers..

The use of morphogenic genes appears to contribute positively to the *Cuscuta* transformation protocol. We observed an enhancement in transformation efficiency with *Cuscuta GRF-GIF*, a phenomenon similar to previously reported findings in wheat and citrus transformation (Debernardi *et al*., 2020). In *Agrobacterium*-mediated *Cuscuta* transformation, *CcGRF-GIF* showed minimal impact on the overall presence/absence ratios of GUS transformation efficiency (Figure 8A). However, the number of GUS segments larger than 2mm was significantly increased in *CcGRF-GIF-GUS* infected *Cuscuta* compared to *Cuscuta* segments that had been transformed with *GUS* alone (Figure 8B). This suggests that *CcGRF-GIF* may not significantly influence *Agrobacterium*-mediated transformation but could enhance the regeneration of transformed tissue, as previously reported (Debernardi *et al*., 2020). Further investigation is needed regarding the utilization of *CcGRF-GIF* with *Cuscuta* transformation. In contrast, using *CcWUS-GUS* in transformations led to lower presence/absence ratios of GUS transformation efficiency than *GUS* alone or *CcGRF-GIF*-*GUS* (Figure 8A). We hypothesize that, similar to previous studies, the constitutive *CcWUS2* expression by the 35S promoter may interfere with the *Cuscuta* development genes, resulting in the decreased transformation efficiency (Rashid *et al*., 2007; Lowe *et al*., 2016; Hoerster *et al*., 2020; Kadri *et al*., 2021).

Observing the interaction of RUBY expressing *Cuscuta* and its hosts suggests that the RUBY reporter system serves not only as a selection marker for transformation, but also as an indicator of events during parasitism at the molecular level. We were able to monitor the invasion progress of the haustorium and observe what appears to be movement of RUBY pigment from the transformed parasite into host cells in the area around developed *Cuscuta* searching hyphae (Figure 6H, K). We do not think this is an artefact of sectioning as the RUBY diffusion was not observed during the early invasion stage (asterisk, Figure 6G). This finding is intriguing given the ability of *Cuscuta* to exchange not only small molecules, but also mobile mRNAs, small RNAs and proteins with its host plants (Kim *et al*., 2014; Park *et al*., 2022)(Shahid *et al*., 2018). Unfortunately, the current data do not allow confident identification of the mechanism of pigment movement. Potential explanations for the movement could include: 1) translation of mobile RNA, wherein mRNAs of betalain synthesis enzymes move from *Cuscuta* to the host stem and are translated near the haustoria; 2) mobile proteins from *Cuscuta* to host, involving the export of three betalain synthesis enzymes from *Cuscuta* that produced betalain in the host stem; or 3) the movement of betalain pigment itself from *Cuscuta* to the host. The last of these options would seem to be the simplest, but betalain is normally sequestered in vacuoles and is not translocated (Chen *et al*., 2017), and *C. campestris* growing on beet does not take up pigment from its host (Park and Westwood, personal observations). Fortunately, we now have the tools to validate these hypotheses. It would be interesting to generate transgenic *Cuscuta* expressing a fluorescence reporter gene fused with endoplasmic reticulum (ER) targeting peptide, such as ER-YFP (Sager *et al*., 2020), which produces a non-mobile product that would provide a definitive marker of parasite cells. Further research is needed, but regardless of the details, the ability to generate transgenic *Cuscuta* as described here will enable many more experiments, and much greater understanding, of *Cuscuta* biology.

## Acknowledgement

This work was supported by the USDA-AFRI-2023-67013-39896 to SP and JW, CAFNR-JOY, University of Missouri to SP, Research Council, University of Missouri to SP and NSF award IOS-1645027 to JW. We thank Duy Trinh who made the *CcGRF-GIF* construction. Grammarly generated responses to the following AI prompts: Prompts created by Grammarly-“Improve it”.

## Author contributions

SP, HL, and JW conceived the ideas and designed methodology. SA, AM, VB, HG, and SP conducted lab-work and analyzed the data. SA, AM, SP, and JW led the writing of the manuscript. All authors contributed critically to the drafts and gave final approval for publication.

## Data availability

The datasets used and/or analyzed during the current study are available from the corresponding author upon request.

## Supplementary

**Supplementary Figure S1.**
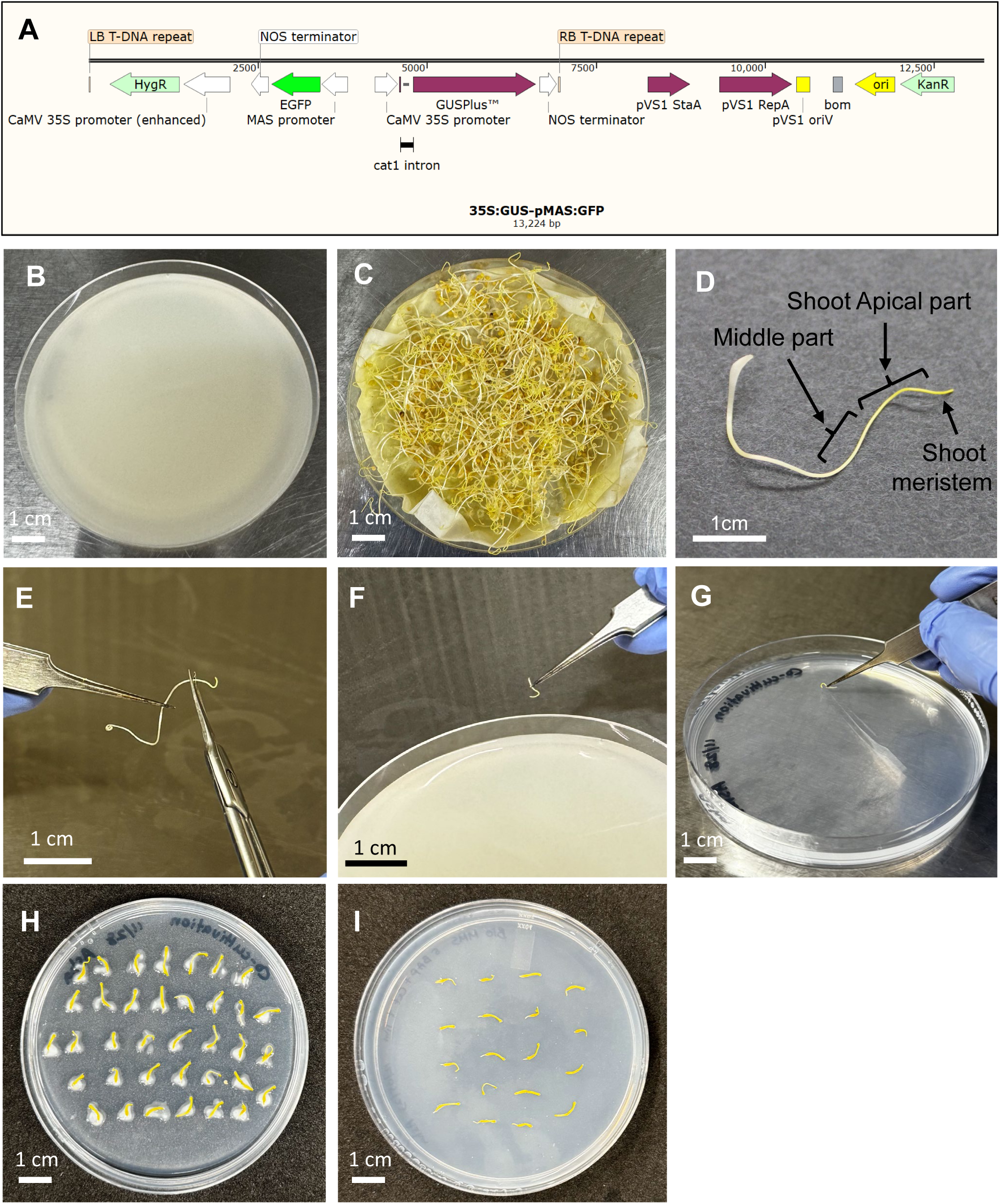
Process of *Agrobacterium*-mediated *Cuscuta* transformation with the first protocol. The vector map of *35S:GUS-pMAS:GFP* used for the transformation (A). Solid creamy bacteria culture (B) and three days old *Cuscuta* seedlings (C) and were prepared on the day of inoculation. To obtain a shoot apical part (D), the shoot meristem and middle part were excised using scissors (E). The excised shoot apical part was then immersed in the creamy culture of *A. tumefaciens* EHA105-*35S:GUS-pMAS:GFP* (F). Explants were placed on the MMS media with 3mg/L NAA and 5mg/L BAP for co-cultivation (G). An excessive *Agrobacteria* were cultivated from infected tissue seven days after inoculation on the co-cultivation media (H). Explants were incubated on MMS with different Auxin and Cytokinin concentrations (I).

**Supplementary Figure S2.**
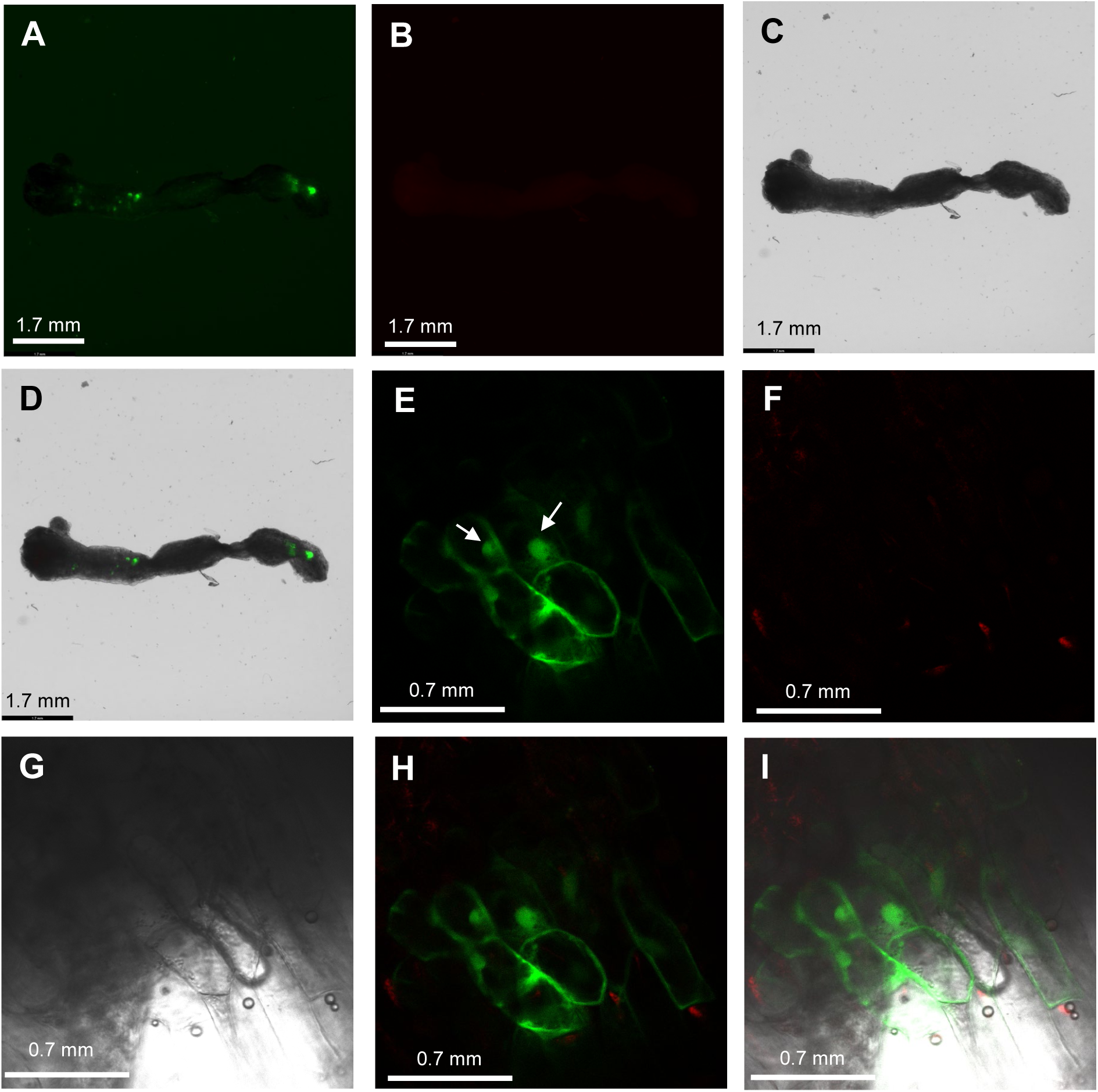
GFP, RFP, and bright field images were captured from approximately 1-month-old explants using a stereomicroscope and a confocal microscope. Leica Thunder imager model organism stereomicroscope system detected GFP (A), RFP (B), and bright field (C) images and then merged these three images (D). Leica SP8 spectral confocal microscope detected GFP (E), RFP (F), and bright field (G) and then merged GFP and RFP (H) and GFP, RFP, and bright field images (I).

**Supplementary Figure S3.**
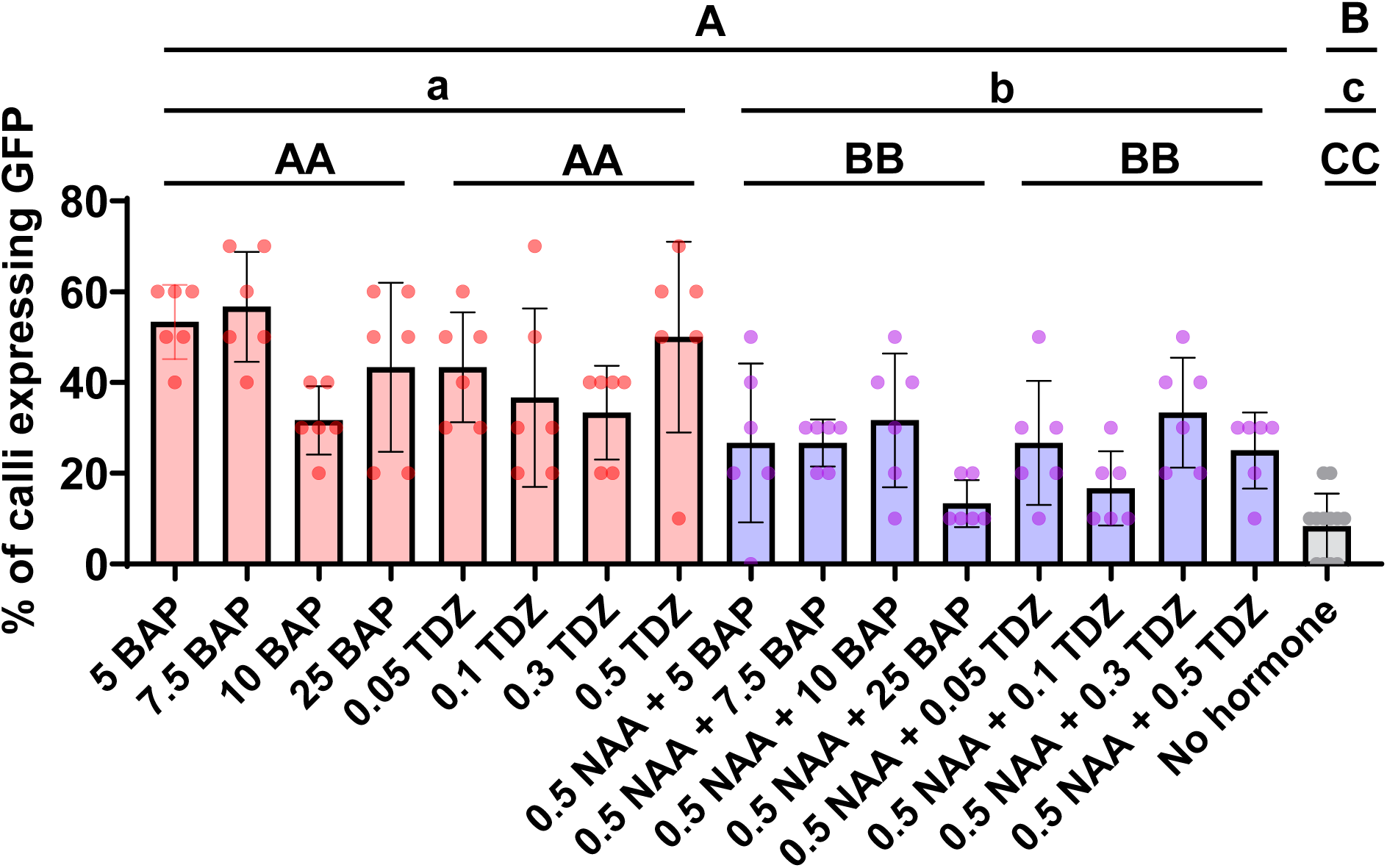
Effect of cytokinins and auxin on GFP positive calli formation after inoculation with *Agrobacterium tumefaciens* EHA105 containing *35S:GUS-pMAS:GFP*. Different types and concentrations of cytokinins (BAP and TDZ) and auxin (NAA) were tested. Six independent biological replicates were performed. Data are presented as the mean ± standard deviation. Groups with different letters vary significantly (two-way ANOVA-Tukey test, P,0.05).

**Supplementary Figure S4.**
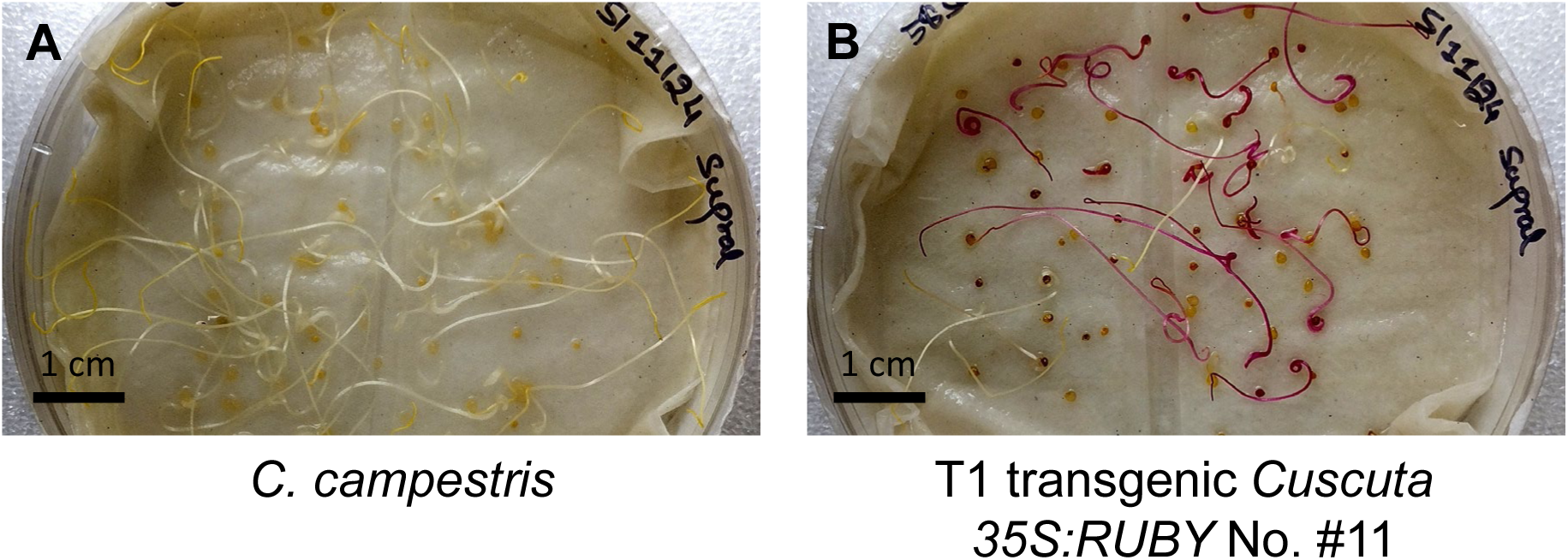
Phenotype of T1 transgenic *Cuscuta 35S:RUBY* No. #11. Three-day-old seedlings of wild-type *C. campestris* (A) and 3-day-old T1 transgenic *Cuscuta 35S:RUBY* No. #11 (B). Scarified WT and transgenic seeds were incubated at 4°C overnight, followed by two days in a 28°C incubator without light, and then one day under long-day dark/light conditions before being imaged.

**Supplementary Figure S5.**
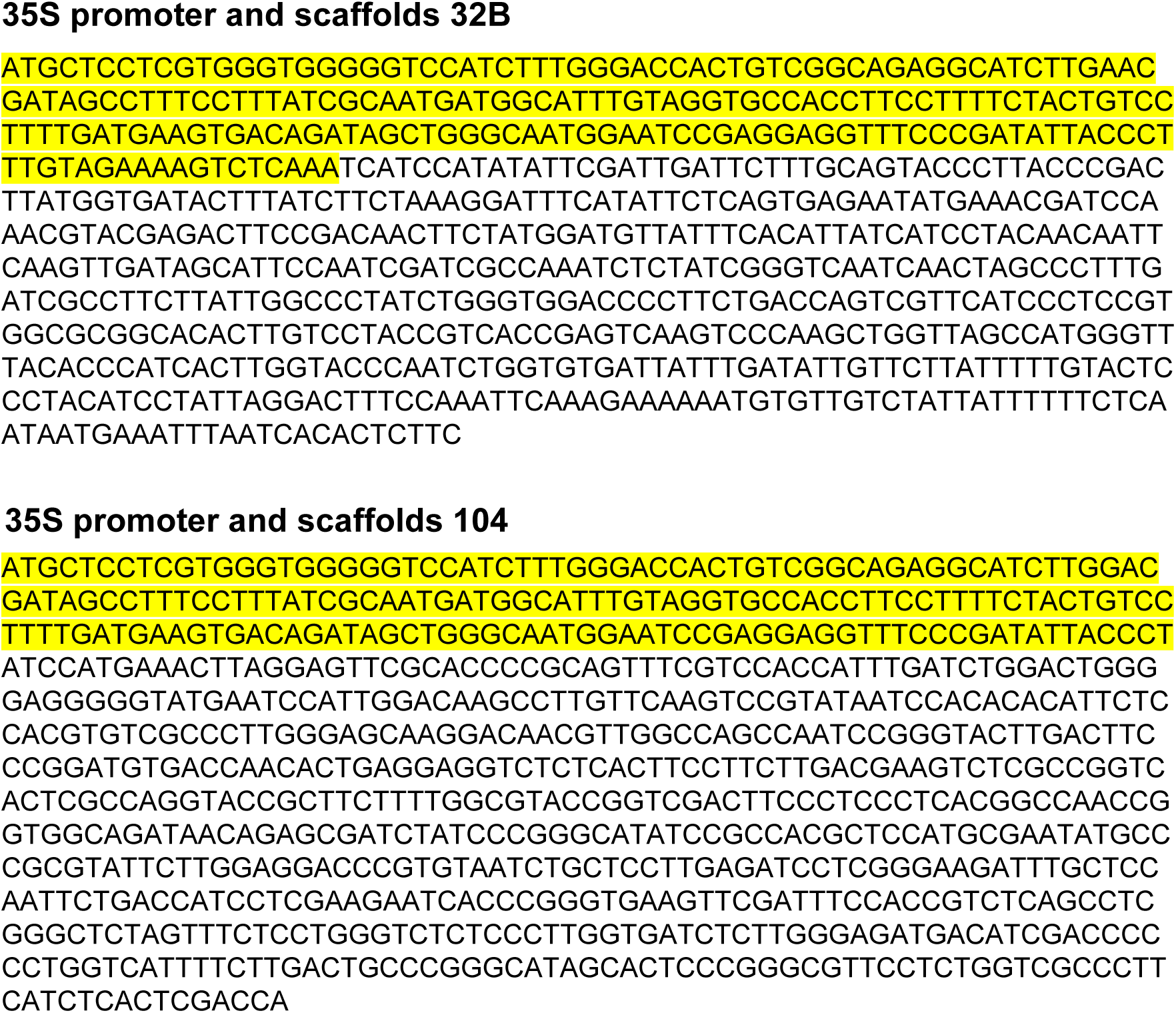
Results of TAIL-PCRs to confirm genomic integration. Sequences are details of the diagram in Figure 5B. The *35S:RUBY* vector sequences (yellow highlights) are linked to *C. campestris* genome scaffolds 32B and 104.

**Supplementary Figure S6.**
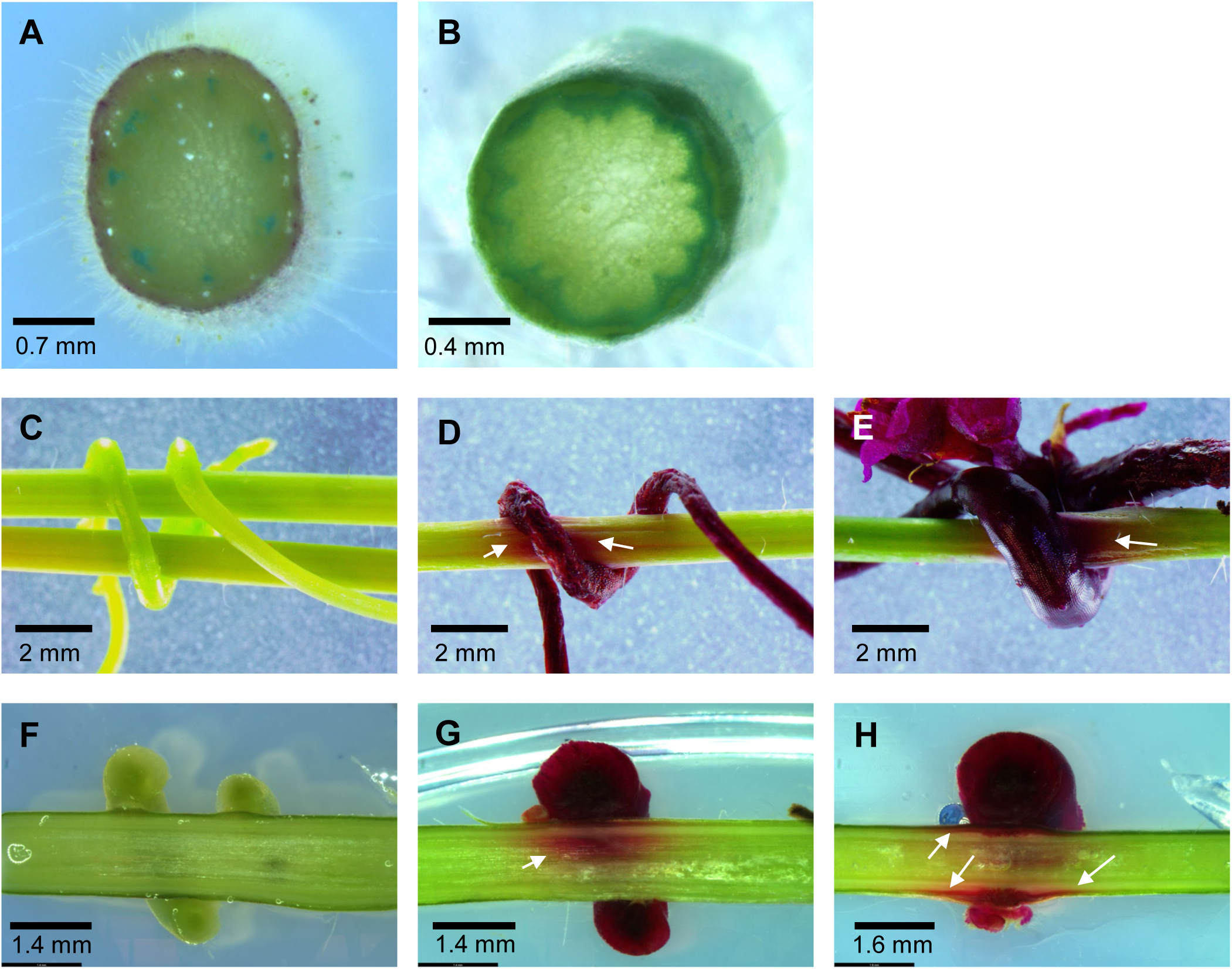
Images from sectioned tomato and *Arabidopsis* stems with WT *Cuscuta* or RUBY *Cuscuta*. Rutgers tomato (A) and *Arabidopsis* Col-0 (B) without *Cuscuta* were sectioned and stained by toluidine blue. WT *Cuscuta* growing on *Arabidopsis* Col-0 (C) and RUBY *Cuscuta* growing on *Arabidopsis* Col-0 (D and E) were vertically sectioned (F-H), respectively. White arrows indicate the RUBY pigment on *Arabidopsis* stems.

**Supplementary Figure S7.**
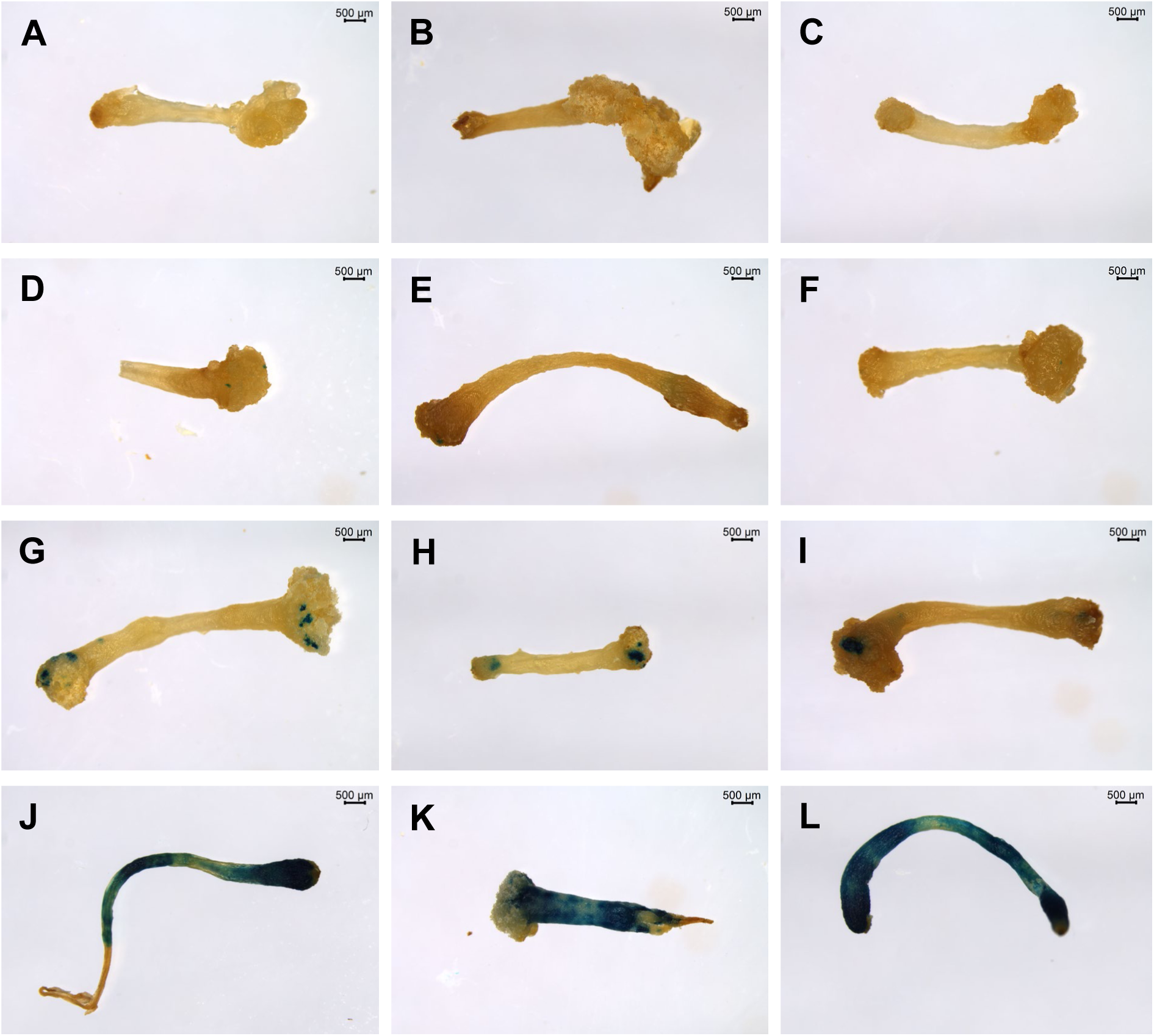
Various sizes of GUS expression sectors in transformed tissues. Following the GUS staining assay, the segments were classified into four groups: No GUS (A-C), less than 0.2 mm GUS (D-F), between 0.2 to 2 mm GUS (G-I), and larger than 2 mm GUS (J-L). The bars represent a scale of 500 µm.

