## Supporting tables for "*Agrobacterium*-mediated *Cuscuta campestris* transformation as a tool for understanding plant-plant interactions"

**Supplementary Table 1.** Media compositions for the first and second protocols.

| **Modified MS Media (MMS) for Co-Cultivation Media (CCM) and Callus Induction Media (CIM)** | |
|---|---|
| **Component** | **Amount** |
| $NH_4NO_3$ (Phytotech labs) | 300 mg/L |
| $KNO_3$ (Fisher) | 800 mg/L |
| $CaCl_2.2H_2O$ (Sigma) | 250 mg/L |
| $MgSO_4.7H_2O$ (Fisher) | 260 mg/L |
| $KH_2PO_4$ (Fisher) | 120 mg/L |
| $H_3BO_3$ (Fisher) | 6.2 mg/L |
| $MnSO_4.4H_2O$ (Sigma) | 22.3 mg/L |
| $ZnSO_4.7H_2O$ (Sigma) | 8.6 mg/L |
| KI (Sigma) | 0.83 mg/L |
| $NaMO_4.2H_2O$ (Sigma) | 0.25 mg/L |
| $CuSO_4.5H_2O$ (Sigma) | 0.025 mg/L |
| $CoCl_2.6H_2O$ (Sigma) | 0.025 mg/L |
| Myo-inositol (Sigma) | 100 mg/L |
| Nicotinic acid (Sigma) | 1 mg/L |
| Pyridoxine hydrochloride (Sigma) | 1 mg/L |

| | |
|---|---|
| Thiamine hydrochloride (Sigma) | 10 mg/L |
| Glucose (Sigma) | 3% (w/v) |
| Phyto-agar (PlantMedia) | 8 g/L |
| pH = 5.7-5.8 | |
| **Co-Cultivation Media (CCM)** | |
| **Component** | **Amount** |
| MMS media | |
| NAA (Sigma) | 3 mg/L |
| BAP (Sigma) | 1 mg/L |
| **Callus Induction Media (CIM)** | |
| **Component** | **Amount** |
| MMS media | |
| Cefotaxime (GoldBio) | 250 mg/L |
| Different amounts of NAA, BAP and TDZ for the first protocol* | |
| NAA (Sigma) | 0, 0.5 (mg/L) |
| BAP (Sigma) | 5, 7.5, 10, 25 (mg/L) |
| TDZ (Sigma) | 0.05, 0.1, 0.3, 0.5 (mg/L) |
| For second protocol | |

| | |
|---|---|
| BAP (Sigma) | 5 mg/L |

* Different amounts of plant growth regulators were applied as described in the methods section.

**Supplementary Table 2.** Media compositions for the third protocol.

| **Co-cultivation Media (CCM)** | |
|---|---|
| Same as the first and second protocols (Table 1) | |
| **Callus Induction Media (CIM)** | |
| **Component** | **Amount** |
| MMS media in Table 1 | |
| BAP (Sigma) | 5 mg/L |
| Cefotaxime (GoldBio) | 250 mg/L |
| Timentin (GoldBio) | 100 mg/L |
| **MS modified basal salt mixture (SKU: 30630204-2, PlantMedia) with B5 vitamins and sugar** | |
| Component | Amount |
| $NH_4NO_3$ | 825 mg/L |
| $KNO_3$ | 950 mg/L |
| $CaCl_2.2H_2O$ | 166.1 mg/L |
| $MgSO_4.7H_2O$ | 180.7 mg/L |
| $KH_2PO_4$ | 170 mg/L |

| | |
|---|---|
| $H_3BO_3$ | 6.2 mg/L |
| $MnSO_4.H_2O$ | 16.9 mg/L |
| $ZnSO_4.7H_2O$ | 8.6 mg/L |
| KI | 0.83 mg/L |
| $NaMO_4.2H_2O$ | 0.25 mg/L |
| $CuSO_4.5H_2O$ | 0.025 mg/L |
| $CoCl_2.6H_2O$ | 0.025 mg/L |
| $FeSO_4.7H_2O$ | 27.8 mg/L |
| $Na_2EDTA.2\ H_2O$ | 37.26 mg/L |
| Myo-inositol (Sigma) | 100 mg/L |
| Nicotinic acid (Sigma) | 1 mg/L |
| Pyridoxine hydrochloride (Sigma) | 1 mg/L |
| Thiamine hydrochloride (Sigma) | 10 mg/L |
| Glucose (Fisher) | 3% (w/v) |
| Phyto agar (PlantMedia) | 8 g/L |
| pH = 5.7-5.8 | |

| **Shoot Induction Media (SIM)** | |
|---|---|
| **Component** | **Amount** |
| MS modified basal salt mixture (SKU: 30630204-2, PlantMedia), B5 vitamins, sugar | |
| BAP (Sigma) | 5 mg/L |
| Cefotaxime (GoldBio) | 250 mg/L |
| Timentin (GoldBio) | 100 mg/L |
| **Shoot Elongation Media (SEM)** | |
| **Component** | **Amount** |
| MS modified basal salt mixture (SKU: 30630204-2, PlantMedia), B5 vitamins, sugar | |
| $CuSO_4.5H_2O$ (Sigma) | 5 mg/L |
| Cefotaxime (GoldBio) | 250 mg/L |
| Timentin (GoldBio) | 100 mg/L |

**Supplementary Table 3.** TAIL-PCR conditions.

| Primary TAIL-PCR | Secondary TAIL-PCR |
|---|---|
| 95C – 3 min | 95C – 3 min |
| 5 cycles | 5 cycles |
| 95C – 30s | 95C – 10s |
| 62C – 1 min | 64C – 1min |
| 72C – 2:30 min | 72C – 1:30min |
| 2 cycles | 15 cycles |
| 95C – 30s | 95C – 10s |
| 25C – 3min (50% ramp) | 64C – 1 min |
| 72C – 2:30 min (32% ramp) | 72C – 1:30 min |
| | 95C -10s |
| | 64 – 1 min |
| 15 cycles | 72C – 1:30 min |
| 95C – 10s | 95C -10s |
| 68 – 1min | 44C – 1 min |
| 72C – 2:30 min | 72C – 1:30 min |
| 95C -10s | |
| 68 – 1min | 5 cycles |
| 72C – 2:30 min | 95C – 10s |
| 95C – 10s | 44C – 1 min |
| 44C – 1 min | 72C – 2min |
| 72C – 2:30 min | |
| | 72C – 5min |
| 72C – 5min | 4C 2min |
| 4C – 2min | |

**Supplementary Table 4**. Primers used for TAIL PCR.

| | Name | Sequence 5'-3' |
|---|---|---|
| **T-DNA specific primers** | RB1-RUBY | GTGTCCTCTCCAAATGAAATGAACTTCCTTATATAGAG |
| | RB2-RUBY | CCTTTGGTCTTCTGAGACTGTATCTTTGATATTCTTG |
| **Arbitrary primers** | AD1 | NGTCGASWGANAWGAA |
| | AD2 | TGWGNAGSANCASAGA |
| | AD3 | AGWGNAGWANCAWAGG |
| | AD6 | WGTGNAGWANCANAGA |

**Supplementary Table 5**. Primers used for cloning.

| | Forward primers | Reverse primers |
|---|---|---|
| *CcGRFGIF* | ATGATGAGCAGTGGAAGAGGCAGG | tTTGATTCCCGCCCGCCGATC |
| *CCWUS2* | ATGGAGCCTCAACACTACTATC | tGGCGAAATGGTGAGACGAC |

**Supplementary Table 6.** Transformation efficiency on different callus induction media

| NAA (mg/L) | BAP (mg/L) | TDZ (mg/L) | Plate no | Number GFP expressing explants | Total explants per plate | Frequency of transformation (%) | Average (%) |
|---|---|---|---|---|---|---|---|
| 0 | 5 | 0 | 1 | 6 | 10 | 60 | 53.3 |
| 0 | 5 | 0 | 2 | 6 | 10 | 60 | |
| 0 | 5 | 0 | 3 | 5 | 10 | 50 | |
| 0 | 5 | 0 | 4 | 4 | 10 | 40 | |
| 0 | 5 | 0 | 5 | 5 | 10 | 50 | |
| 0 | 5 | 0 | 6 | 6 | 10 | 60 | |
| 0 | 7.5 | 0 | 1 | 5 | 10 | 50 | 56.7 |
| 0 | 7.5 | 0 | 2 | 5 | 10 | 50 | |
| 0 | 7.5 | 0 | 3 | 7 | 10 | 70 | |
| 0 | 7.5 | 0 | 4 | 6 | 10 | 60 | |
| 0 | 7.5 | 0 | 5 | 4 | 10 | 40 | |
| 0 | 7.5 | 0 | 6 | 7 | 10 | 70 | |
| 0 | 10 | 0 | 1 | 3 | 10 | 30 | 31.7 |
| 0 | 10 | 0 | 2 | 3 | 10 | 30 | |
| 0 | 10 | 0 | 3 | 4 | 10 | 40 | |
| 0 | 10 | 0 | 4 | 4 | 10 | 40 | |
| 0 | 10 | 0 | 5 | 2 | 10 | 20 | |
| 0 | 10 | 0 | 6 | 3 | 10 | 30 | |
| 0 | 25 | 0 | 1 | 6 | 10 | 60 | 43.3 |
| 0 | 25 | 0 | 2 | 5 | 10 | 50 | |
| 0 | 25 | 0 | 3 | 6 | 10 | 60 | |
| 0 | 25 | 0 | 4 | 5 | 10 | 50 | |
| 0 | 25 | 0 | 5 | 2 | 10 | 20 | |
| 0 | 25 | 0 | 6 | 2 | 10 | 20 | |
| 0 | 0 | 0.05 | 1 | 5 | 10 | 50 | 43.3 |

| | | | | | | | |
|---|---|---|---|---|---|---|---|
| 0 | 0 | 0.05 | 2 | 3 | 10 | 30 | |
| 0 | 0 | 0.05 | 3 | 4 | 10 | 40 | |
| 0 | 0 | 0.05 | 4 | 6 | 10 | 60 | |
| 0 | 0 | 0.05 | 5 | 3 | 10 | 30 | |
| 0 | 0 | 0.05 | 6 | 5 | 10 | 50 | |
| 0 | 0 | 0.1 | 1 | 7 | 10 | 70 | 36.7 |
| 0 | 0 | 0.1 | 2 | 2 | 10 | 20 | |
| 0 | 0 | 0.1 | 3 | 3 | 10 | 30 | |
| 0 | 0 | 0.1 | 4 | 5 | 10 | 50 | |
| 0 | 0 | 0.1 | 5 | 3 | 10 | 30 | |
| 0 | 0 | 0.1 | 6 | 2 | 10 | 20 | |
| 0 | 0 | 0.3 | 1 | 2 | 10 | 20 | 33.3 |
| 0 | 0 | 0.3 | 2 | 4 | 10 | 40 | |
| 0 | 0 | 0.3 | 3 | 2 | 10 | 20 | |
| 0 | 0 | 0.3 | 4 | 4 | 10 | 40 | |
| 0 | 0 | 0.3 | 5 | 4 | 10 | 40 | |
| 0 | 0 | 0.3 | 6 | 4 | 10 | 40 | |
| 0 | 0 | 0.5 | 1 | 6 | 10 | 60 | 50.0 |
| 0 | 0 | 0.5 | 2 | 5 | 10 | 50 | |
| 0 | 0 | 0.5 | 3 | 6 | 10 | 60 | |
| 0 | 0 | 0.5 | 4 | 7 | 10 | 70 | |
| 0 | 0 | 0.5 | 5 | 5 | 10 | 50 | |
| 0 | 0 | 0.5 | 6 | 1 | 10 | 10 | |
| 0.5 | 5 | 0 | 1 | 5 | 10 | 50 | 26.7 |
| 0.5 | 5 | 0 | 2 | 2 | 10 | 20 | |
| 0.5 | 5 | 0 | 3 | 4 | 10 | 40 | |
| 0.5 | 5 | 0 | 4 | 3 | 10 | 30 | |
| 0.5 | 5 | 0 | 5 | 2 | 10 | 20 | |
| 0.5 | 5 | 0 | 6 | 0 | 10 | 0 | |
| 0.5 | 7.5 | 0 | 1 | 2 | 10 | 20 | 26.7 |

| | | | | | | | |
|---|---|---|---|---|---|---|---|
| 0.5 | 7.5 | 0 | 2 | 2 | 10 | 20 | |
| 0.5 | 7.5 | 0 | 3 | 3 | 10 | 30 | |
| 0.5 | 7.5 | 0 | 4 | 3 | 10 | 30 | |
| 0.5 | 7.5 | 0 | 5 | 3 | 10 | 30 | |
| 0.5 | 7.5 | 0 | 6 | 3 | 10 | 30 | |
| 0.5 | 10 | 0 | 1 | 1 | 10 | 10 | 31.7 |
| 0.5 | 10 | 0 | 2 | 4 | 10 | 40 | |
| 0.5 | 10 | 0 | 3 | 5 | 10 | 50 | |
| 0.5 | 10 | 0 | 4 | 4 | 10 | 40 | |
| 0.5 | 10 | 0 | 5 | 2 | 10 | 20 | |
| 0.5 | 10 | 0 | 6 | 3 | 10 | 30 | |
| 0.5 | 25 | 0 | 1 | 1 | 10 | 10 | 13.3 |
| 0.5 | 25 | 0 | 2 | 1 | 10 | 10 | |
| 0.5 | 25 | 0 | 3 | 2 | 10 | 20 | |
| 0.5 | 25 | 0 | 4 | 1 | 10 | 10 | |
| 0.5 | 25 | 0 | 5 | 2 | 10 | 20 | |
| 0.5 | 25 | 0 | 6 | 1 | 10 | 10 | |
| 0.5 | 0 | 0.05 | 1 | 5 | 10 | 50 | 26.7 |
| 0.5 | 0 | 0.05 | 2 | 2 | 10 | 20 | |
| 0.5 | 0 | 0.05 | 3 | 3 | 10 | 30 | |
| 0.5 | 0 | 0.05 | 4 | 2 | 10 | 20 | |
| 0.5 | 0 | 0.05 | 5 | 3 | 10 | 30 | |
| 0.5 | 0 | 0.05 | 6 | 1 | 10 | 10 | |
| 0.5 | 0 | 0.1 | 1 | 1 | 10 | 10 | 16.7 |
| 0.5 | 0 | 0.1 | 2 | 2 | 10 | 20 | |
| 0.5 | 0 | 0.1 | 3 | 1 | 10 | 10 | |
| 0.5 | 0 | 0.1 | 4 | 3 | 10 | 30 | |
| 0.5 | 0 | 0.1 | 5 | 2 | 10 | 20 | |
| 0.5 | 0 | 0.1 | 6 | 1 | 10 | 10 | |
| 0.5 | 0 | 0.3 | 1 | 2 | 10 | 20 | 33.3 |

| | | | | | | | |
|---|---|---|---|---|---|---|---|
| 0.5 | 0 | 0.3 | 2 | 5 | 10 | 50 | |
| 0.5 | 0 | 0.3 | 3 | 3 | 10 | 30 | |
| 0.5 | 0 | 0.3 | 4 | 4 | 10 | 40 | |
| 0.5 | 0 | 0.3 | 5 | 2 | 10 | 20 | |
| 0.5 | 0 | 0.3 | 6 | 4 | 10 | 40 | |
| 0.5 | 0 | 0.5 | 1 | 3 | 10 | 30 | |
| 0.5 | 0 | 0.5 | 2 | 1 | 10 | 10 | |
| 0.5 | 0 | 0.5 | 3 | 2 | 10 | 20 | 25.0 |
| 0.5 | 0 | 0.5 | 4 | 3 | 10 | 30 | |
| 0.5 | 0 | 0.5 | 5 | 3 | 10 | 30 | |
| 0.5 | 0 | 0.5 | 6 | 3 | 10 | 30 | |
| 0 | 0 | 0 | 1 | 2 | 10 | 20 | |
| 0 | 0 | 0 | 2 | 2 | 10 | 20 | |
| 0 | 0 | 0 | 3 | 1 | 10 | 10 | |
| 0 | 0 | 0 | 4 | 0 | 10 | 0 | |
| 0 | 0 | 0 | 5 | 1 | 10 | 10 | |
| 0 | 0 | 0 | 6 | 1 | 10 | 10 | 8.3 |
| 0 | 0 | 0 | 1 | 0 | 10 | 0 | |
| 0 | 0 | 0 | 2 | 1 | 10 | 10 | |
| 0 | 0 | 0 | 3 | 0 | 10 | 0 | |
| 0 | 0 | 0 | 4 | 0 | 10 | 0 | |
| 0 | 0 | 0 | 5 | 1 | 10 | 10 | |
| 0 | 0 | 0 | 6 | 1 | 10 | 10 | |

**Supplementary Table 7.** Different transformation efficiency between the cut and the bulk chop method. DAI is days after inoculation.

| **Wounding method** | **Incubation dates** | **Total explants per plate** | **Replicates** | | | | | **Average number of red spots in a plate** | **Average efficiency (%)** |
|---|---|---|---|---|---|---|---|---|---|
| | | | **plate 1** | **plate 2** | **plate 3** | **plate 4** | **plate 5** | | |
| Cut | 20 DAI | 10 | 10 | 10 | 10 | 10 | 10 | 10 | 100 |
| Chop | 20 DAI | 20 | 15 | 17 | 10 | 9 | 14 | 13 | 65 |

**Supplemental Table 8.** Transformation efficiency with RUBY reporter. The number of segments expressing RUBY was evaluated from 4 to 45 days after Agro inoculation (DAI). Sixteen independent biological repeats were performed. Inoculums from two hundred seeds were deviled into two CIM, each containing 50 explants per plate.

| No. of reps. | 5 DAI (%) | 15 DAI (%) | 25 DAI (%) | 35 DAI (%) | 45 DAI (%) |
|---|---|---|---|---|---|
| 1 | 100 | 88.0 | 82.0 | 78.0 | 68.0 |
| 2 | 100 | 88.5 | 80.8 | 76.9 | 69.2 |
| 3 | 100 | 94.6 | 80.4 | 73.2 | 60.7 |
| 4 | 100 | 92.3 | 82.7 | 69.2 | 65.4 |
| 5 | 100 | 98.0 | 84.0 | 74.0 | 50.0 |
| 6 | 100 | 100.0 | 94.0 | 82.0 | 60.0 |
| 7 | 100 | 88.1 | 78.0 | 74.6 | 45.8 |
| 8 | 100 | 98.1 | 94.2 | 78.8 | 57.7 |
| 9 | 100 | 94.0 | 94.0 | 80.0 | 80.0 |
| 10 | 100 | 90.0 | 90.0 | 74.0 | 74.0 |
| 11 | 100 | 84.0 | 80.0 | 74.0 | 62.0 |
| 12 | 100 | 96.0 | 94.0 | 74.0 | 72.0 |
| 13 | 100 | 94.7 | 86.0 | 84.2 | 52.6 |
| 14 | 100 | 96.2 | 90.4 | 88.5 | 65.4 |
| 15 | 100 | 96.2 | 84.6 | 73.1 | 53.8 |
| 16 | 100 | 92.9 | 80.4 | 73.2 | 57.1 |
| Average | 100 | 93.2 | 86.0 | 76.7 | 62.1 |

**Supplemental Table 9.** Transgenic RUBY *Cuscuta*. The number of RUBY calli growing on SEM, the number of RUBY calli producing shoots, and the number of RUBY *Cuscuta* coiled on a host were counted and quantified. Eight independent biological repeats were performed, each replicate contains 50 explants per plate.

| No. of reps. | % of RUBY calli growing on SEM | % of RUBY calli producing shoots | % of RUBY *Cuscuta* coiled on a host |
|---|---|---|---|
| 1 | 24.0 | 10.0 | 8.0 |
| 2 | 14.0 | 4.0 | 4.0 |
| 3 | 12.0 | 4.0 | 2.0 |
| 4 | 24.0 | 8.0 | 4.0 |
| 5 | 18.0 | 8.0 | 2.0 |
| 6 | 6.0 | 0.0 | 0.0 |
| 7 | 10.0 | 0.0 | 0.0 |
| 8 | 20.0 | 8.0 | 2.0 |
| Average | 16.0 | 5.3 | 2.8 |

**Supplementary Table 10.** Germination efficiency and segregation of RUBY phenotype in different lines of the *35S:RUBY* transgenic *Cuscuta* and wild type.

| Line | Replicates | Total number of seeds | Germinating seedlings | Average germination efficiency | STDEV of germination efficiency | Number of RUBY seedlings | Number of non-colored seedlings | The Ratio of RUBY/ non-RUBY seedlings | Average RUBY Ratio: non-RUBY seedlings | STDEV of RUBY Ratio: non-RUBY seedlings |
|---|---|---|---|---|---|---|---|---|---|---|
| Wild Type *Cuscuta campestris* | R1 | 10 | 60.0% | 70.0% | ± 17.3% | 0 | 6 | 0 | 0 | 0 |
| | R2 | 10 | 90.0% | | | 0 | 9 | 0 | | |
| | R3 | 10 | 60.0% | | | 0 | 6 | 0 | | |
| Seeds from T0 *35S:RUBY* No. #11 | R1 | 10 | 90.0% | 86.7% | ± 5.8% | 6 | 3 | 2 : 1 | 2.33 : 1 | ± 0.58 |
| | R2 | 10 | 80.0% | | | 6 | 2 | 3 : 1 | | |
| | R3 | 10 | 90.0% | | | 6 | 3 | 2 : 1 | | |
